# Species boundaries to the limit: validating species delimitation methods is critical to avoid taxonomic inflation in the genomic era

**DOI:** 10.1101/2022.12.28.522089

**Authors:** Bernat Burriel-Carranza, Maria Estarellas, Gabriel Riaño, Adrián Talavera, Héctor Tejero-Cicuéndez, Johannes Els, Salvador Carranza

## Abstract

With the advent of molecular phylogenetics, the number of taxonomic studies unveiling and describing cryptic diversity has greatly increased. However, speciation between cryptic lineages is often defined without evaluating population structure or gene flow, which can lead to false claims of species status and, subsequently, taxonomic inflation. In the present study we focus on the intriguing case of the Arabian gecko *Trachydactylus hajarensis* (Squamata: Gekkonidae), a species for which cryptic diversity has been previously reported. We generated mitochondrial data (12S rDNA) and genome-wide SNP data (ddRADseq) for 52 specimens to determine phylogenomic relationships, population structure and gene flow within this species. Then, we applied species delimitation methods (SDMs) to evaluate several competing species hypotheses through the Multispecies Coalescent model. Results show that *T. hajarensis* is comprised by three well-defined lineages, two of them in the Hajar Mountains of eastern Arabia, and one in Masirah Island, in the southeastern coast of Oman. Even though high levels of past introgression and strong mitonuclear discordances were found, current gene flow is scarce with clear boundaries between populations and shallow levels of admixture in the contact zone between lineages. Surprisingly, species tree topology differed between methods and when different individuals were used in downsampled datasets. Conventional SDMs supported up to three putative new species within the group. However, after species validation with the genealogical divergence index (*gdi*), none of the putative species held. Overall, this study highlights the importance of sample choice, integrative analyses, and validation methods to not incur into taxonomic inflation, providing a set of already available tools to assess and validate population structure, gene flow, and SDMs before describing new species.

## 1. Introduction

Species are one of the fundamental units in biology. However, with more than 24 different definitions, the species concept is still a topic of debate, and sometimes dispute, in the scientific community (Mayden, 1997). Many authors agree that species are separately evolving metapopulations emerging from a speciation process (Fišer, Robinson, & Malard, 2018) but, since speciation is intrinsically a gradual mechanism, discriminating populations from species remains a challenge. The inference of speciation events is achieved by attending to different lines of evidence such as reproductive isolation, morphological differentiation or monophyly (De Queiroz, 2007; Fišer et al., 2018). However, the advent of molecular phylogenetics and genomics has presented a new paradigm, reshaping the tree of life as we knew it. We have seen that phenotypic variation or the absence of gene flow alone do not always suffice to delimit species and, in many cases, species present signals of ancestral or current hybridization (Ivanov, Lee, & Mutanen, 2018). Moreover, the increase of molecular species delimitation studies in the last decade has revealed that cryptic diversity is much more common than previously thought (Chattopadhyay et al., 2016; Vences et al., 2022; Vilaça et al., 2021).

Cryptic species are entities that have undergone a speciation process but remain morphologically identical (Chan et al., 2020). Owing to the generalized use of molecular methods in most systematic studies, it has been shown that cryptic species are widely spread across most animal phyla (Derycke et al., 2008; Fennessy et al., 2016; Riaño et al., 2022; Vences et al., 2022). Some of the mechanisms that can lead to a lack of phenotypical variation between divergent lineages include early divergence or niche conservatism (Fišer et al., 2018), among others. Moreover, the description of cryptic diversity is shortening the Linnean shortfall and is of paramount importance for conservation efforts (Walters, Cannizzaro, Trujillo, & Berg, 2021), since many species that were once thought to be homogeneous and widely distributed, actually represent complexes of cryptic species with sometimes endangered micro-endemic entities (Garcia-Porta, Simó-Riudalbas, Robinson, & Carranza, 2017). However, the lack of a specific threshold to distinguish separate evolutionary lineages from populations of the same species raises another challenging question: where do we stop? The advent of next generation sequencing (NGS) and the development of new species delimitation methods (SDMs) have led to an outburst of taxonomic studies, especially in the case of non-model organisms (Ivanov et al., 2018). The amount of data that can be obtained through these techniques has massively surpassed the multi-locus approach and, in several cases, signals of population structure can be interpreted as species status with conventional SDMs, even when there is gene flow (Sukumaran & Knowles, 2017). This might result in over-splitting and taxonomic inflation, an issue especially problematic for conservation management, where limited resources have to be prioritized to the most endangered species (Isaac, Mallet, & Mace, 2004).

Nowadays, genomic SDMs are at the forefront of an ongoing debate in the scientific community (Jackson et al., 2017; Leaché et al., 2019; Sukumaran & Knowles, 2017). Within those, the Multispecies Coalescent (MSC) model (Rannala & Yang, 2003), extensively used with genomic data from closely related species, has been shown to capture population splits rather than species divergences (Leaché et al., 2019). This is especially problematic in allopatric populations where a lack of gene flow does not confirm pre- or postzygotic barriers but rather results from geographic isolation. To account for this, Jackson et al. (2017) proposed a heuristic criterion for species delimitation based on a genealogical divergence index (*gdi*) which afterwards was implemented in a Bayesian parameter estimation under the MSC model (Leaché et al., 2019). With this index, two lineages can be identified either as two distinct species or as a single one, but it also includes a range of indecision that reflects the arbitrary nature of the species definition (Leaché et al., 2019).

In the past decades, molecular phylogenetics and the application of SDMs in traditionally neglected arid regions have proven that in such areas there are still high levels of undescribed diversity and many examples of cryptic species (Bray, Alagaili, & Bennett, 2014; Carranza et al. 2016; Main, van Vuuren, Tilbury, & Tolley, 2022; Simó-Riudalbas et al., 2017). Within the Arabian Peninsula, the Hajar Mountains rise as one of its most biodiverse regions, with high levels of reptile diversity and endemicity (Burriel-Carranza, Els, & Carranza, 2022; Carranza, Els, & Burriel-Carranza, 2021; Šmíd et al., 2021). Its high and complex topography and the relatively low annual mean temperatures offer a spectrum of diverse niches which have already been the scenery of the origin of cryptic diversity (Garcia-Porta et al., 2017; Simó-Riudalbas et al., 2017, 2018; Tamar, Mitsi, & Carranza, 2019). Therefore, it is a well-suited ecological system to explore the nuances of the MSC model together with the new heuristic *gdi* with allopatric and early divergent species.

The ground-dwelling Arabian gecko *Trachydactylus hajarensis* (Arnold, 1980) is a species endemic to the Hajar Mountains and to Masirah Island, a small island situated 20 km off the eastern coast of Oman, but almost 200 km away from the closest continental *T. hajarensis* (Figure 1). Even though previous data suggest a natural colonization of the island instead of a human-mediated translocation (de Pous et al., 2016), the provenance of this island’s population still remains unclear. Previous studies on this South Arabian endemic genus of only two species were carried out with multi-locus assemblies and most of the analyses were based on mitochondrial sequences alone (de Pous et al., 2016). Based on those results, *T. hajarensis* was suggested to be a species complex with at least three allopatric lineages: one in the Western and Central Hajars, a second in the Central and Eastern Hajars, and another one in the easternmost side of the Hajar Mountains and in Masirah Island (de Pous et al., 2016). However, mitochondrial evolutionary history is not always linked to its nuclear counterpart, with common cases of mito-nuclear discordances across phyla (Marshall, Chambers, Matz, & Hillis, 2021; Shults et al., 2022). This stresses the need of revisiting this group’s systematics with NGS techniques, which recover large portions or the complete nuclear genome and are key to determine whether there are mito-nuclear discordances, providing a comprehensive perspective of the evolutionary history of a species.

**Figure 1.**
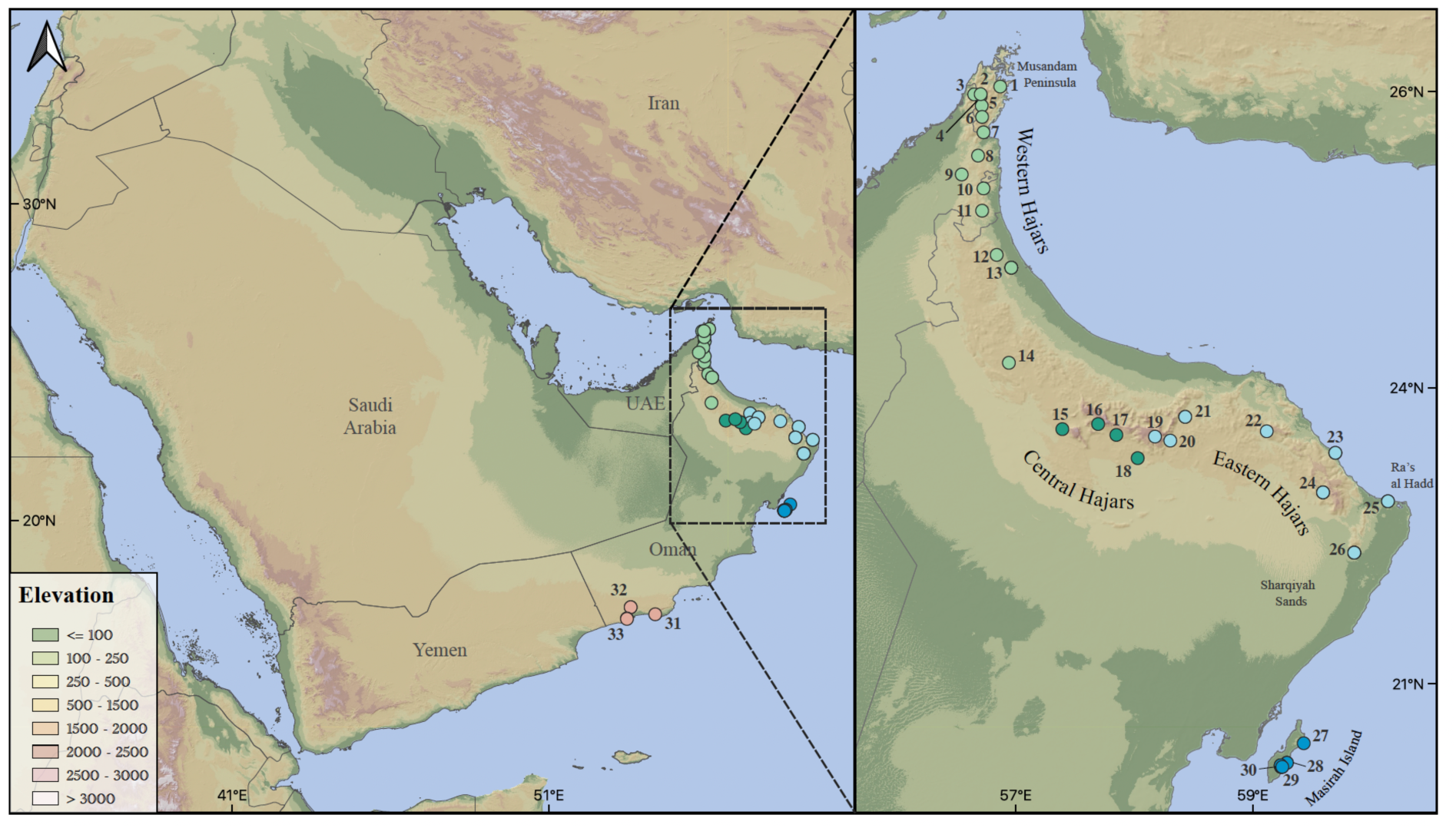
Topographic map of the study area showing the 33 localities of *Trachydactylus* sampled for this study (see Table S1 for correspondece between the 33 localities and the 53 sampled specimens); colors represent *T. spatalurus* (pink) and *T. hajarensis* Western (light green), Central (dark green), Eastern (light blue) and Masirah Island (dark blue) lineages according to our ddRADseq genomic analyses.

In the present work, we generate *de novo* genomic reduced representations of *T. hajarensis* to revisit the systematics of the group. We reconstruct the nuclear and mitochondrial phylogeny of *T. hajarensis* together with its sister species *T. spatalurus*, generate and test a series of species hypotheses applying the heuristic *gdi* and delimit the population structure and evolutionary history of this species, with a special focus on the colonization of Masirah Island.

## 2. Materials & Methods

### 2.1. Sampling

A total of 52 individuals of the genus *Trachydactylus* from 33 different localitites were included in this study (Figure 1; Table S1). Specimens were collected between 2005 and 2017 along all the known distribution range of *T. hajarensis*, containing representatives of all the Hajar Mountains’ and Masirah Island’s lineages. We also included four specimens of *T. spatalurus* from Dhofar, South Oman, that were used as outgroup (Table S1).

### 2.2. Mitochondrial analyses

We sequenced 35 specimens for the 12S rDNA (*12S*) and 15 additional samples were downloaded from GenBank. DNA extraction was done following the protocol in MacManes (2013) and PCR amplification conditions and primers used were the same as described in Metallinou et al. (2015). PCR products were purified and Sanger sequenced by Macrogen Inc. to obtain a 523 bp fragment of the mitochondrial gene *12S*. Sequences were aligned with Geneious 2021.1.1 (Biomatters Ltd.) and a Bayesian Inference (BI) phylogeny was reconstructed in BEAST2 v.2.6.4 (Bouckaert et al., 2019). We calibrated the deepest node in our phylogeny (the split between *T. hajarensis* and *T. spatalurus*) by extracting the mean height from a recently published squamate phylogeny (Tejero-Cicuéndez et al., 2022) and applying a normal distribution encompassing the 95% HPD intervals. We selected a HKY+G+X model with four gamma categories, base frequencies were estimated, and a relaxed clock LogNormal was used with a Calibrated Yule process tree prior. We conducted three independent runs of 10^8^ generations sampling every 10,000 generations. Convergence was checked with Tracer v.1.5 (Rambaut & Drummond, 2013), a 40% burnin was applied and trees were summarized with TreeAnnotator v.2.6.4 (Bouckaert et al., 2019). Then, we objectively identified and delimited deep mitochondrial lineages using the general mixed Yule-coalescent model (GMYC; Fujisawa & Barraclough, 2013; Pons et al., 2006) implemented in the R package ‘splits’ (Ezard, Fujisawa, & Barraclough, 2021).

### 2.3. Genomic DNA sequencing and processing

Genomic libraries were produced following Peterson’s et al. (2012) protocol for double-digest restriction site-associated DNA (ddRADseq) for 52 specimens. In short, we double-digested 500 ng of genomic DNA, using a pair of rare and common restriction enzymes (Sbf1 and Msp1, respectively). The resulting fragments were ligated with barcoded Illumina adapters. Fragments were then size-selected for a range between 415 – 515 bp and sequenced on an Illumina NextSeq 500, for 75 bp single-end reads.

Raw Illumina reads were processed using Ipyrad v.0.9.62 (Eaton & Overcast, 2020) discarding sites with Phred score < 33, reads with more than three missing sites, consensus sequences with less than six reads, excessive heterozygous sites (more than three), or more than two haplotypes. After testing several configurations, both filtered reads and consensus sequences were clustered and aligned using an 89% clustering threshold. We set the minimum number of samples per locus to four to retrieve the maximum number of loci possible for post-processing filtering. Demultiplexed filtered reads for each individual can be found at Dryad (will be added in the published version).

Following recommendations from O’Leary et al. (2018), we applied an iterative filtering to identify and remove low quality samples and loci. We used Radiator (Gosselin, Lamothe, Devloo-Delva, & Grewe, 2017), Plink2 (Chang et al., 2015) and VCFr (Knaus & Grünwald, 2017) implemented in a custom script (https://github.com/BernatBurriel/Post_processing_filtering) to filter iteratively and alternatively SNP datasets. Values of missing data allowance ranged from 98% to 78% of missing genotype call rate and missing data per individual, decreasing 2% between iterations. Furthermore, we applied a hard threshold of missing genotype call rate depending on the dataset (Table S2), we removed non-biallelic SNPs, applied a minor allele frequency filter (maf < 0.05) and removed monomorphic sites.

Dataset types varied between analyses but can be summarized into *loci* and *SNP* datasets: *loci* datasets were generated with ipyrad after removing all individuals that did not pass the previously explained filters and retaining only loci that were at least in 60% of all specimens. *SNP* datasets contained either concatenated SNPs or putatively unlinked SNPs. In the latter only the SNP with the highest read depth of each locus was chosen. For further specifications in each dataset, refer to Table S2.

### 2.4. Population structure

We used *dataset 1* (Table S2) to inferred the population ancestry of each individual with ADMIXTURE v.1.3.0 (Alexander & Lange, 2011; Alexander, Novembre, & Lange, 2009). Ancestral populations ranged from K=1 to K=10, with 15 replicates for each K, and the best K value inferred after 15 cross-validation rounds. Then, we used the same dataset to perform a Principal Component Analysis (PCA) with Plink v2.00a2.3 (Chang et al., 2015). Finally, we used *dataset 2* (Table S2) for fineRADstructure (Malinsky, Trucchi, Lawson, & Falush, 2018) to further analyze *T. hajarensis’* population structure. This analysis unravels different levels of structure within and between populations and its robustness to missing data is optimal for non-model organisms (Malinsky, Trucchi, Lawson, & Falush, 2018). Results from all the former analyses were visualized with R v.4.2.1 (R Core Team, 2021).

### 2.5. Phylogenomic reconstructions

We conducted the phylogenomic anlayses using ML and BI on *dataset 3* (Table S2), including 5,219 loci and 47 individuals. With a concatenated dataset of all loci, we generated ML reconstructions with RAxML-ng v.1.0.2 (Kozlov et al., 2019) with a GTR+G model, a total of 100 starting trees (50 random and 50 parsimony) and 1,000 bootstrap replicates to estimate branch support. We also generated individual gene trees for each locus with IQ-TREE (Nguyen, Schmidt, von Haeseler, & Minh, 2015) with the best model obtained from ModelFinder (TPM3+F+R2), 100 trees and 1,000 ultrafast bootstraps. Then we generated a consensus tree by summarizing all trees with Astral v.5.7.8 (Zhang, Sayyari, & Mirarab, 2017).

We also estimated a time calibrated tree with BI implemented in BEAST2 v.2.6.4 (Bouckaert et al., 2019) applying the same priors and specifications as in the mitochondrial BI approach (see methods section 2.2). In addition, we reconstructed with *dataset 3* the relationships of *T. hajarensis* using unrooted phylogenetic networks implemented in SplitsTree v.4.18.3 (Huson & Bryant, 2006) with the Neighbor-Net algorithm.

### 2.6. Coalescent-based Species Trees

Based on the population structure (Figure 2) and phylogenomic reconstruction (Figure 3) analyses, we identified up to four well-defined monophyletic groups within *T. hajarensis*. These groups can be geographically divided into Western Hajars lineage, Central Hajars lineage, Eastern hajars lineage and Masirah Island lineage (Figure 1). We estimated a species tree of the aforementioned lineages together with *T. spatalurus* to evaluate the evolutionary relationships between these groups in the Multispecies Coalescent framework. The resulting tree was then used as a guide tree for species delimitation methods (see methods section 2.8). Since species delimitation and species tree inference tend to be computationally demanding, it is common practice to downsample datasets to one or two specimens per lineage (e.g. Tonzo, Papadopoulou, & Ortego, 2019). To test the effect of taxon choice when producing reduced datasets, we generated four datasets with different specimen configurations that did not present any signals of admixture between lineages (see Table S2 for further details); *T. spatalurus* was used to root the phylogenetic trees: i) *dataset 4 SNAPP/BPP*: two specimens with the highest coverage from Western, Central, Eastern, and Masirah Island lineages and two specimens of *T. spatalurus*; ii) *dataset 5 SNAPP/BPP*: two specimens from Western, Central, Eastern, and Masirah Island lineages and two specimens of *T. spatalurus*; Eastern lineage specimens selected from the closest geographic region to the Masirah Island lineage (CN10775-26 and CN10791-26); iii) *dataset 6 SNAPP/BPP*: two specimens from Western, Central, Eastern, and Masirah Island lineages and two specimens of *T. spatalurus*; Eastern lineage specimens selected from the farthest geographic region to the Masirah Island lineage (S7150-22 and CN686-23); vi) *dataset 7 SNAPP/BPP:* four specimens from Western, Central, Eastern, and Masirah Island lineages and four specimens of *T. spatalurus*; Eastern lineage specimens selected from all its geographic range (S7150-22, S7161-24, CN4226-26 and CN10775-26). For further information on the selected specimens and dataset specifications refer to Tables S1 and S2.

**Figure 2.**
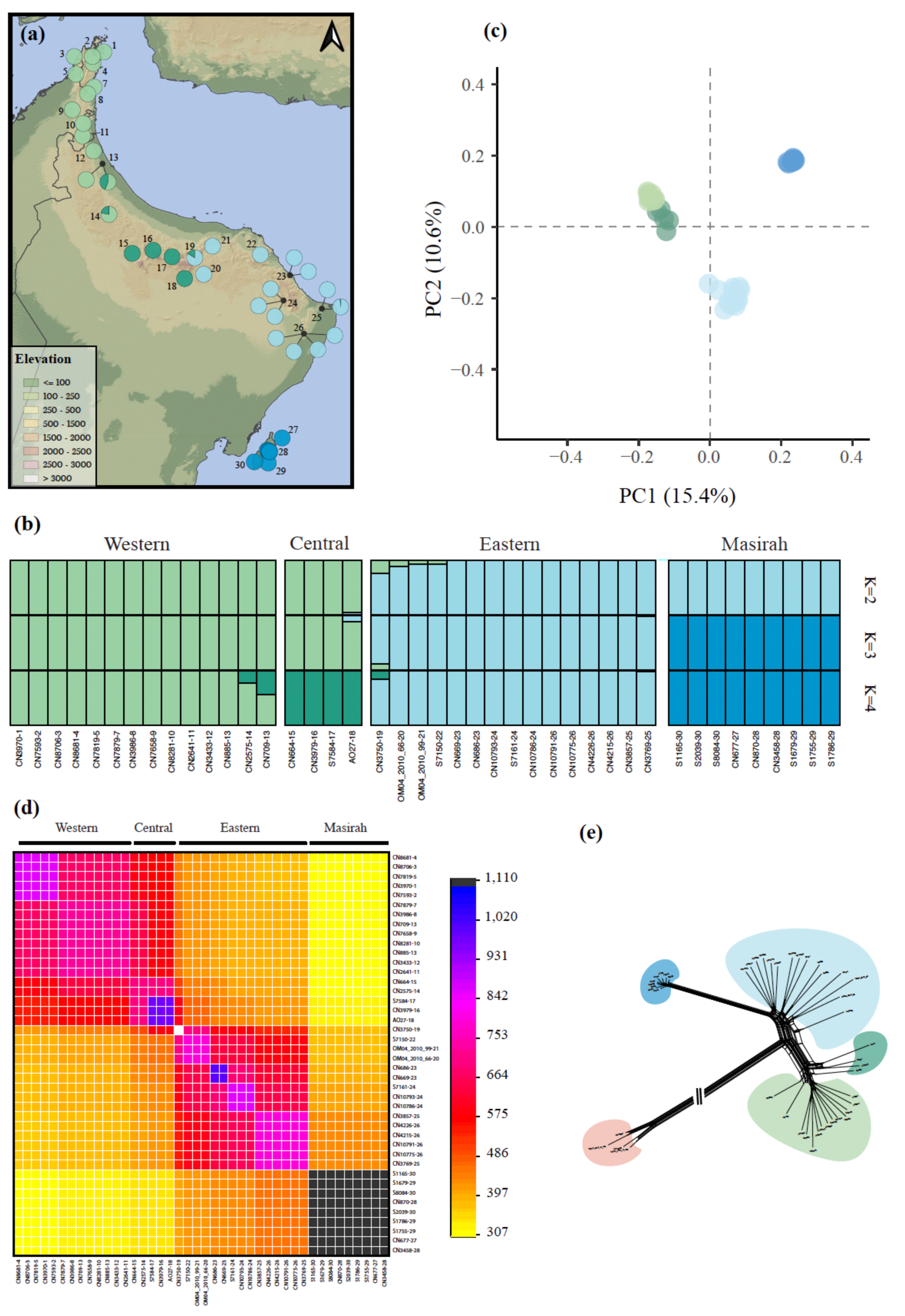
Population structure analyses of *T. hajarensis.* **a**: Geographic distribution of the different individuals analyzed, with pies representing the percentage of admixture for K=3; **b**: individual assignments of ADMIXTURE nuclear clusters (K=2-4); Numbers after last dash correspond to locality number from Figure 1 and Table S1; **c**: Principal Component Analysis (PCA), with points representing the different individuals colored by taxa; **d**: Co-ancestry matrix generated in fineRADstructure, indicating pairwise genetic similarity between all *T. hajarensis* specimens. To the right, legend shows the number of shared alleles between individuals. Darker colors representent higher inter-individual co-ancestry; **e:** Phylogenomic network constructed with the Neighbor-Net algorithm in SplitsTree. *T. spatalurus* (pink); *T. hajarensis* Western (light green), Central (dark green), Eastern (light blue) and Masirah Island (dark blue) lineages. Datasets contain 2,428 unlinked SNPs (**a**,**b** & **c**; *dataset 1*), 30,526 loci (**d**; *dataset 2*) and 5,219 loci (**e**; *dataset 3*). See table S2 for further specifications on each dataset.

**Figure 3.**
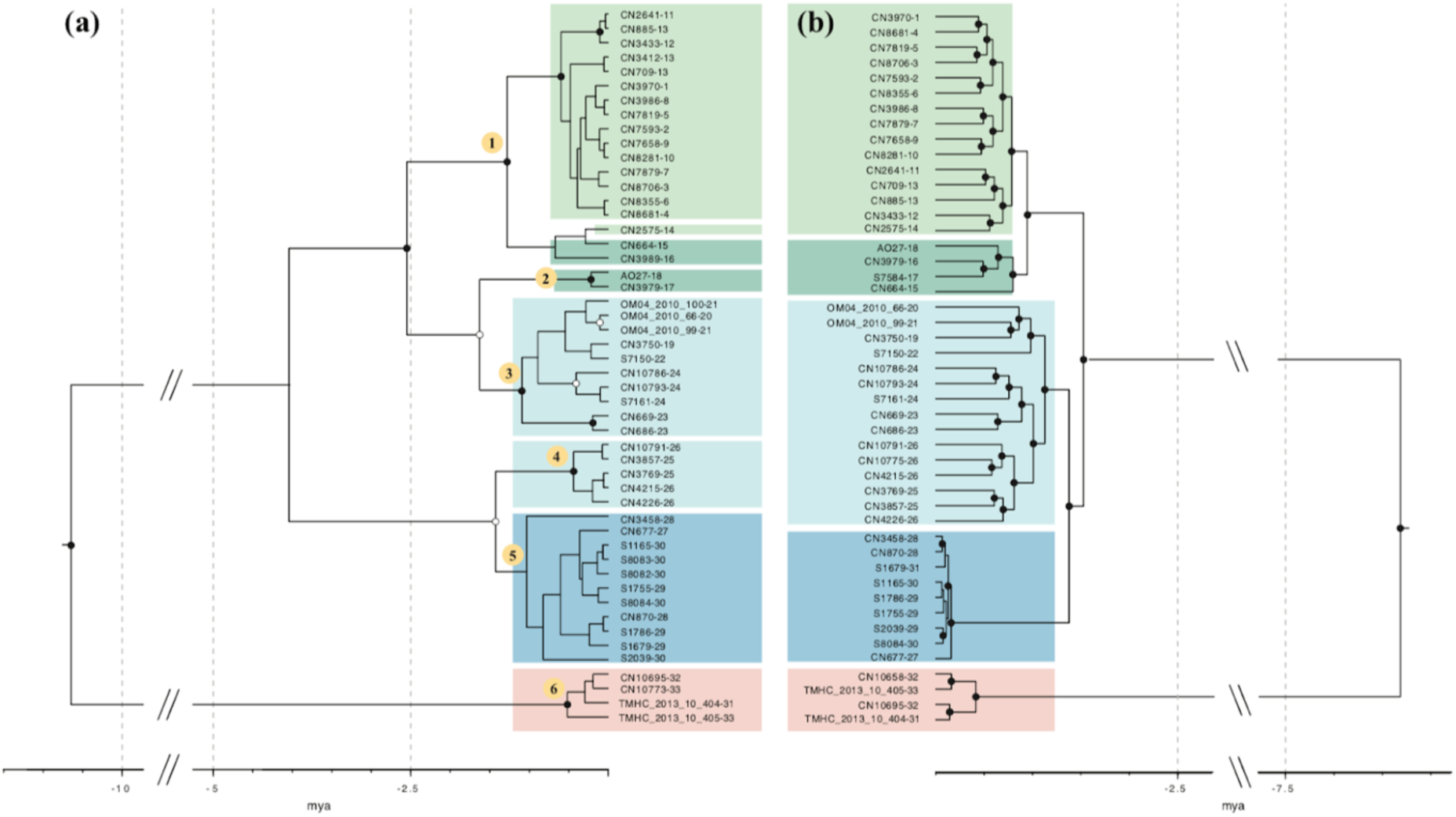
Bayesian inference time calibrated trees. **a**: Phylogenetic tree based on the *12S* mitochondrial gene. Numbers in yellow circles correspond to GMYC species level assignment; **b**: Phylogenomic tree inferred with a concatenated dataset of 5,219 nuclear loci. Posterior probability (pp) above 0.95 and pp > 0.85 are shown with black and white dots at each node respectively; Numbers after last dash correspond to locality number in Figure 1 and Table S1; *T. spatalurus* (pink), *T. hajarensis* Western (light green), Central (dark green), Eastern (light blue) and Masirah Island (dark blue) lineages.

Time-calibrated species trees for each of the four datasets above were inferred with SNAPP v.1.5.2 (Bryant et al. 2012) twice. First, we generated a time-calibrated species tree with SNAPP using the ‘snapp_prep.rb’ script (https://github.com/mmatschiner/tutorials). We dated the deepest node in the phylogeny as suggested by Stange et al. (2018) with a normal distribution from the mean age extracted from Tejero-Cicuéndez et al. (2022). Mutation rates (*u* & *v*) were fixed to 1, and a uniform distribution was set for the population mutation rate theta (*θ*) with default boundaries (0-1,000) and was constrained to be identical on all branches. The latter prior is assumed by the script ‘snapp_prep.rb’ to decrease the computational load of the analysis (Stange et al., 2018). We also repeated the same analysis without linking population mutation rates and applying specific Yule (*λ*) and Theta priors (*θ*; see Coalescent-based Species Delimitation below). In both approaches we ran 4 independent runs of 3,000,000 generations, sampling every 50 generations. Convergence between runs and stationarity was checked with Tracer v.1.7 (Rambaut & Drummond, 2013). Posterior distributions were combined with LogCombiner v.2.6.3, discarding 30% of the posterior trees as burn-in and a maximum clade credibility tree was obtained calculating median heights in TreeAnnotator v.2.6.3 (BEAST2 v.2.6.4; Bouckaert et al., 2019).

In addition, we generated species tree estimations for *dataset 4 BPP* to *dataset 7 BPP* (Table S2) with BPP A01 analysis (Flouri, Jiao, Rannala, & Yang, 2018). We followed the pipeline of Huang (2018) (https://github.com/airbugs/Dynastes_delimitation) to estimate *θ* and τ priors (a=3, b=0.061 and a=3, b=0.131 respectively), and produced a dataset where all loci were present in at least one individual of each lineage. Then, we implemented three independent runs of 500,000 generations sampling every 10 generations after a burn-in of 50,000.

Finally, we generated a ML species tree with *dataset* 3 (Table S2). We followed the same procedure in IQ-TREE (Nguyen et al., 2015) as in Methods 2.5 but when summarizing all the trees with Astral v.5.7.8 (Zhang et al., 2017) we assigned each individual to its group as inferred by ADMIXTURE.

### 2.7. Species tree topology testing

As shown in results section 3.3.2, our coalescent-based species trees recovered two different topologies: the first topology (from now on *Topo1*) separates *T. hajarensis* into two clades: the first conformed by Western and Central Hajars’ lineages, and the other by Eastern Hajars and Masirah Island’s lineages. The second topology (from now on *Topo2*) places all Hajar Mountains’ lineages as a clade sister to Masirah Island’s. Both trees were rooted with *T. spatalurus*. We evaluated both topologies by implementing the recently published mixture across sites and trees (MAST) model (Wong et al., 2022). In short, this program calculates topology weights for a number of given topologies (two in our case) across an alignment, calculates the most supported topology at each site, and returns an overall value for the whole alignment (determined by a topology weight ranging between 0 and 1). We generated a MAST analysis for each polymorphic loci present in each quintet of specimens containing one individual from all *T. hajarensis* lineages (Western, Central, Eastern and Masirah) and one individual of *T. spatalurus*. There were a total of 37,800 quintets which contained, on average, 537 ±141 loci extracted from *dataset 3* (Table S2). Then, we ran MAST independently for each loci (19.35×10^6^ different MAST analyses) giving the aforementioned topologies as input trees and unlinking substitution models, DNA frequencies and gamma model across trees.

Resulting overall tree weights for each loci were extracted, weights not supporting either of the given topologies with a probability above 0.55 were discarded, and the resulting 3.96×10^6^ weights were summarized by specimen and quintet independently. We identified the most supported topology for each specimen by averaging the number of loci between all quintet combinations where a specific specimen was present. When grouping our results by quintet, we implemented a linear model to identify up to which point distance to the putative contact zone (distance to Central Hajars in *Topo1*, and distance to Masirah in *Topo2*) explained the observed shifts in topology weights. We calculated euclidean distances between all Eastern *vs* Masirah Island and Eastern *vs* Central Hajars specimens and accounted for all quintet combinations where Eastern and Masirah/Central specimens remained identical in two different ways: On the one hand, we obtained a single topology weight for each locality by averaging all pseudoreplicates. On the other, we accounted for all pseudoreplicates implementing a nested analysis of variance.

### 2.8. Coalescent-based Species Delimitation

In the present work, we use the General Lineage Species Concept (de Queiroz,2007). This unified species concept considers species as separately evolving meta-population lineages and treats this property as the single requisite for delimiting species. We consider continuously distributed populations that show contemporary gene flow and no other notable forms of divergence to be a metapopulation lineage of the same species (Chan et al., 2020). Other properties, such as phenetic distinguishability, reciprocal monophyly, and pre- and postzygotic reproductive isolation, are not part of this species concept but serve as important line s of evidence relevant to assess the separation of lineages and therefore species status (de Queiroz, 2007).

We designed and tested several coalescent-based species delimitation models by splitting or lumping the previously inferred *T. hajarensis* lineages. Specifically, our species delimitation hypotheses included: *i)* Single-species hypothesis (*H_0_*); *ii*) Two species within *T. hajarensis* (*H_1_*: Western+Central *vs* Masirah + Eastern lineages); *iii*) Three-species hypothesis (*H_2_*: Western + Central *vs* Eastern *vs* Masirah lineages); and *vi*) All lineages as distinct species (*H_3_*: Western *vs* Central *vs* Eastern *vs* Masirah lineages). Species delimitation analyses were tested with Bayes factor delimitation (BFD* with genomic data; Leaché et al. 2014) implemented in BEAST2 v.2.6.4 (Bouckaert et al., 2019), and with BPP A10 analysis (Yang & Rannala, 2010).

Within BFD*, we estimated for each model a species tree with SNAPP v.1.5.2 (Bryant et al., 2012) and conducted a path sampling analysis to estimate and rank marginal likelihoods between models. We then computed Bayes Factors (BF) to determine the best SDM. Since SNAPP is computationally intensive and assumes no gene flow between species, we selected 2 non-admixed individuals from each species hypothesis (*dataset 4 SNAPP*; Table S2). We also included representatives of its sister species *T. spatalurus* to account for *T. hajarensis* being a single species, resulting in a dataset of 10 individuals and 2,147 unlinked SNPs. Mutation rates (*u* & *v*) were fixed to 1. We set the Yule prior (*λ*) to a gamma distribution and while alpha was set to 2, beta was estimated by calculating the expected tree height on the 5,219 loci dataset (maximum observed divergence between any pair of taxa divided by 2). We then used ‘pyule’ (https://github.com/joaks1/pyule) to determine the mean value of lambda and calculated beta accordingly (*λ = α × β*). Theta prior (*θ*) was also set to a gamma distribution and the mean value of *θ* was estimated by averaging all genetic distances within each lineage. Path sampling analyses were run for 20 steps with the following parameters: 500,000 MCMC generations sampling every 1,000, with an alpha of 0.3, 10% burnin and a pre-burnin of 50,000. Stationarity of all runs was checked and each step was run until ESS >= 200.

We implemented further guided species delimitation analyses with BPP v.4.4.1 (Rannala & Yang, 2013; Yang & Rannala, 2010). We used *dataset 4 BPP* which contained the same individuals as in BFD* (Table S2), used the best species tree topology (*Topo1*; see results 3.4) as the guide tree, and processed the dataset as described above (see methods 2.6). Then, we implemented three independent runs of BPP A10 species delimitation (Yang & Rannala, 2010) with 1,000,000 generations, sampling every 10 generations after a burn-in of 100,000.

### 2.9. Species validation

To test the robustness of our inferred species, we calculated the genealogical divergence index (*gdi*) proposed by Jackson et al. (2017), and implemented within BPP by Leaché et al. (2019). The equation for *gdi* is 1 − *e*^−2τAB/θA^, where τ represents the divergence time between species *A* and *B* and θ represents the population size of the species *A*. Therefore, to effectively test species status for two given taxa this index has to be calculated reciprocally between taxa *A* and *B*. According to Jackson et al. (2017), low *gdi* values (*gdi* < 0.2) indicate that *A* and *B* are the same species while high values (*gdi* > 0.7) support a distinct species status of the two taxa. Values in between are considered ambiguous, lacking the support necessary to be classified as distinct species.

We implemented this index to the two competing species tree hypotheses (See results 3.3.2). We ran A00 analyses with *dataset 4 BPP* (Table S2) estimating the parameters on the inferred guide trees (*Topo1* and *Topo2*) to generate the posterior distributions for the most recent species divergences, and then we collapsed those tips and repeated the analysis to test species status for the lumped lineages (see Leaché et al., 2019 for a similar approach). Three independent runs of A00 were performed for 100,000 generations, sampling every five generations after a burn-in of 10,000. We calculated and visualized *gdi* indexes with R v.4.2.1 (R Core Team, 2022).

### 2.10. Introgression analysis

We used the D-Statistics (ABBA-BABA tests) implemented in Dsuite (Malinsky, Matschiner, & Svardal, 2021) to test for past signals of introgression between *T. hajarensis* lineages. To do so, we considered four *T. hajarensis* lineages (as supported by BFD* analysis) and performed the analysis providing both *Topo1* and *Topo2* respectively. In all analyses, *T. spatalurus* was used as outgroup. Finally, we calculated the f-branch statistics to infer the excess of shared derived alleles between lineages.

## 3. Results

### 3.1. ddRADseq data processing

Total reads obtained from sequencing added up to 11.6 x 10^7^ and after applying quality filters more than 97% of the raw reads remained, with an average of 2.1 x 10^6^ reads per individual. Post-processing filtering identified up to five individuals with low coverage levels, which were discarded from subsequent analyses. The number of loci within *loci* datasets ranged between 30,526 to 4,441 loci, and *SNP* datasets contained between 30,096 and 2,428 SNPs, depending on the number of individuals used, and the applied filters for each analysis. See Table S2 for further information on each dataset.

### 3.2. Population structure

We explored the population structure of *T. hajarensis* with *dataset 1* (Table S2), which contained 2,428 unlinked SNPs present in at least 60% of all individuals (missing genotype call rate set to 40). The most likely number of ancestral populations recovered from Admixture was K=3 (cross-validation = 0.43). This configuration geographically segregates *T. hajarensis* into Western + Central Hajars, Eastern Hajars, and Masirah Island, with almost no signal of gene flow between them. Lower k numbers lumped together Masirah and Eastern lineages and higher k numbers split the occidental clade into the Central and Western lineages (Figure 2a & b, Figure S1). Similar results were obtained with the Principal Component Analysis (PCA), which also segregated *T. hajarensis* into the three aforementioned clusters. Interestingly, all continental lineages clustered closer together than to the Masirah Island group (Figure 2c).

To further inspect its population structure, we implemented fineRADstructure to *dataset 2* (Table S2). Results again show a clear structure into three groups, being Masirah Island the cluster sharing the highest number of haplotypes between its specimens (up to 1,110; Figure 2d). Surprisingly, the Eastern lineage showed high levels of coancestry with both Masirah Island and Western + Central lineages, suggesting a shared evolutionary history with both of them. Such coancestry was stronger in specimens that where geographically closer to either Masirah Island (localities 25 and 26) or the Central Hajars (localities 19 to 21). This can be clearly appreciated at the crossroads between the Central and Eastern lineages where we find an individual (CN3750-19; Figure 2d) that almost shares the same coancestry with either Central and Eastern lineages, most likely representing a hybrid between them (Figure 2d). Finally, the phylogenomic network also shows the presence of all three lineages and, even though the Eastern Hajars are more related to Masirah, there are some reticulation events that approximate both Central and Eastern groups (Figure 2e).

### 3.3. Phylogenetic reconstructions

#### 3.3.1. Mitochondrial tree reconstruction

The phylogeny reconstructed with the *12S* mitochondrial gene (Figure 3a) is concordant with the results shown in de Pous et al. (2016) with most of the Hajar Mountain’s lineages forming a monophyletic group (localities 1–24). In this group, there are two main lineages: the first lineage spans from the Musandam Peninsula southwards into the westernmost part of the Central Hajars (localities 1–16), and the second is formed by the specimens inhabiting the Central and most of the Eastern Hajars (localities 17 – 24). However, not all *T. hajarensis* inhabiting the Hajar Mountains cluster together, since the two easternmost localities (localities 25 and 26 in Figure 1) were recovered as sister group to the specimens in Masirah Island (Figure 3a).

#### 3.3.2. Phylogenomic reconstructions and topological discordances

All phylogenomic reconstructions were discordant with the mitochondrial phylogeny and, overall, they show a clear geographical structure with three distinct lineages in the Eastern, Central, and Western Hajars. However, we recovered different topologies regarding the position of Masirah Island’s lineage and the phylogenomic relationships between the aforementioned groups. The concatenation-based BI approach recovered Masirah Island and Eastern Hajars lineages as reciprocally monophyletic (Figure 3b). A very similar topology was obtained with the summarized ML phylogeny (Figure S2) with the exception of the specimen CN3750-19, that was recovered as basal to both Masirah Island and Eastern Hajars lineages, and which most likely represents a hybrid specimen between Central and Eastern lineages (Figure 2d). Finally, concatenation-based ML approaches placed Masirah Island’s lineage within the Eastern lineage (Figure S2). All the above support a Western + Central clade as sister to an Eastern + Masirah clade.

Two species tree topologies were recovered across all species tree reconstruction methods (Figure S3). The first follows the same topology obtained with BI and yields a Western + Central clade sister to an Eastern + Masirah clade (*Topo1*). The second topology places all Hajar Mountains’ *T. hajarensis* as a monophyletic group sister to the Masirah Island lineage (*Topo2*), and was recovered in three SNAPP and one BPP species tree analyses (Figure S3). Both topologies are supported with a posterior probability of 1 in all nodes in at least one analysis (Figure S3).

### 3.4. Species tree topology testing

Between the two competing species tree hypotheses, *Topo1* was consistently recovered as the best topology for almost all specimens and quintets. Only three specimens near the contact zone between Central and Eastern Hajar’s populations presented low to negative number of loci supporting *Topo1* (CN3750-19, OM04_2010_66-20 and OM04_2010_99-21; Figure 4a), and it is noteworthy that in the best Admixture scenario only one of the three specimens (CN3750-19) was recovered as admixed (Figure 2a).

**Figure 4.**
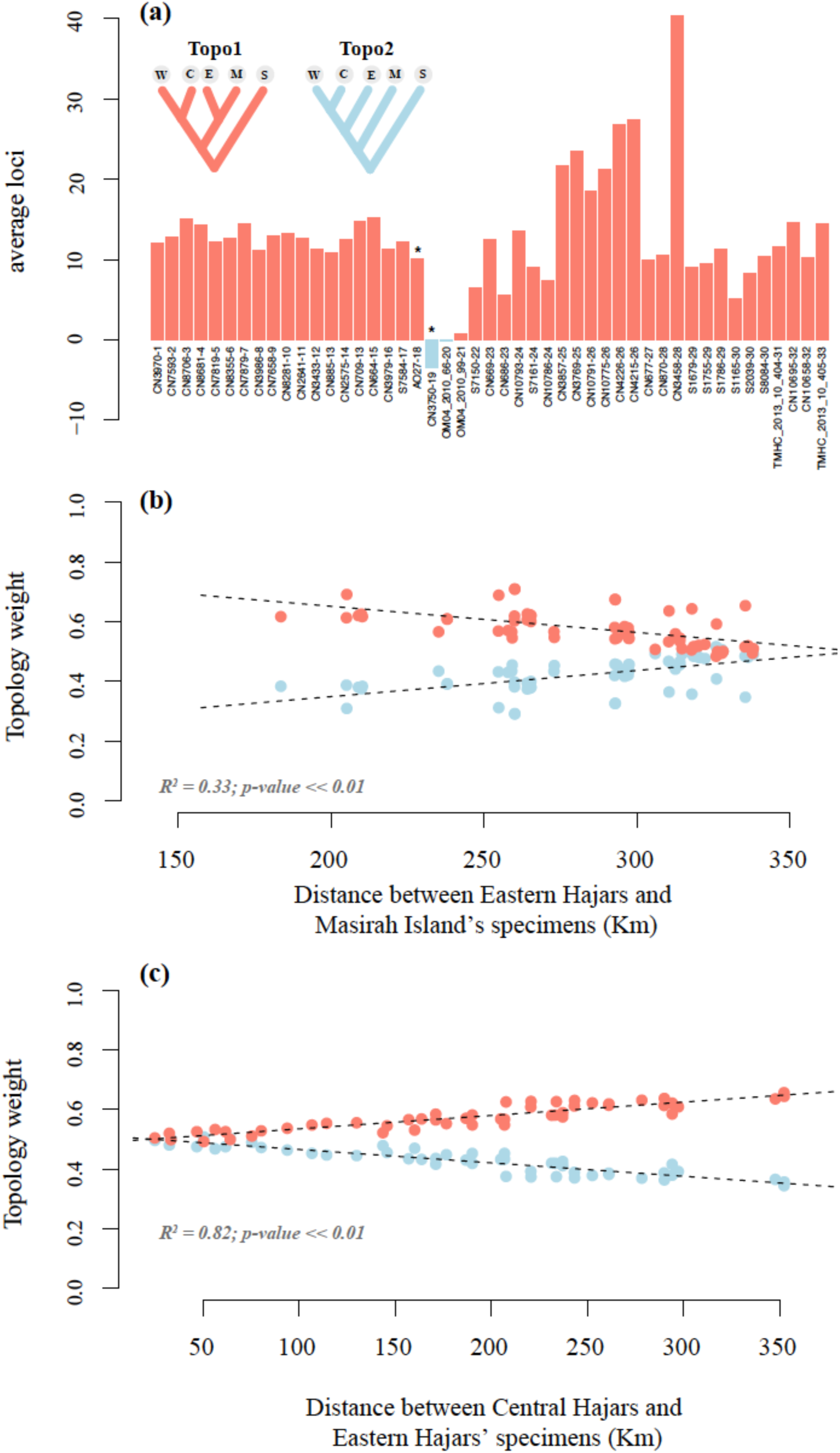
Species tree topology testing. **a:** Number of loci averaged across all quintets that support each topology (shown as *Topo1*-*Topo2*). Numbers after last dash correspond to locality number from Figure 1. Asterisks highlight admixed specimens in Figure 2a. *T. spatalurus* (S), *T. hajarensis* Western (W), Central (C), Eastern (E), and Masirah Island (M) lineages; **b:** Shifts in *Topo1* (salmon) and *Topo2* (blue) topology weights explained by the distance from each Eastern Hajars’ specimen to each Masirah Island’s specimen; **c:** Shifts in *Topo1* (salmon) and *Topo2* (blue) topology weights explained by the distance from each Eastern Hajars’ specimen to each Central Hajars’ individual. Each point corresponds to the averaged topology weight across all quintet combinations with the same Eastern and Masirah specimens (see Figure S4). The sum of topology weights (*Topo1* and *Topo2*) of each MAST analysis always adds up to 1.0, thus plotted regression lines have inverse directions but identical R^2^ and *p*-values.

Splitting our dataset into quintets allowed us to investigate the effects of past gene flow in the observed topology of *T. hajarensis*. When evaluating topology weights through a geographic gradient (either distance between Eastern Hajars and Masirah Island’s specimens in Figure 4b, or distance between Central Hajars and Eastern Hajars’ specimens in Figure 4c) we saw that support for *Topo1* decreased when Eastern Hajars and Masirah Island’s specimens were farther apart (Figure 4b), and when Central and Eastern Hajar’s specimens were closer together (Figure 4c). Both geographic gradients significantly explain the shifts in topology weights (*p*-value << 0.01; Figure 4b,c) and point towards the same direction, a shift from *Topo1* to *Topo2* in the Central/Eastern contact zone. Distance between Central Hajars and Eastern Hajars’ specimens explained much better the observed topology weights (R^2^ = 0.82; Figure 4c) than distance between Eastern Hajars and Masirah Island’s specimens (R^2^ = 0.33; Figure 4b), which is concordant with *Topo1* representing the *true* evolutionary history of *T. hajarensis*. Altogether, results suggest that *Topo2* is the result of a secondary contact between Central and Eastern lineages. The gradual higher support of *Topo2* when Eastern Hajars’ specimens are closer to the Central Hajars instead of a clear-cut shift in the contact zone could be indicative of past dispersals of Central Hajars’ specimens into the Eastern block, leaving a genomic footprint in the nowadays Eastern Hajars’ populations.

Pseudoreplicate or sample choice (different Western Hajars and Central Hajars samples for each distance between Eastern *vs* Masirah Island’s specimens; Figure S4a; different Western Hajars and Masirah Island samples for each distance between Central *vs* Eastern Hajars’ specimens; Figure S4b) was recovered as significant but with less power than distance between Eastern Hajars and the Masirah Island (Table S3) or Central Hajars’ specimens (Table S4). This means that if we downsample our dataset and select far enough Eastern and Central Hajars’ specimens, we will obtain *Topo1* regardless of the other samples in the quintet. However, if we select geographically close Eastern and Central Hajars’ specimens, we can obtain either *Topo1* or *Topo2* in our resulting species tree, depending on the other Western Hajars and Masirah Island samples selected (Figure S4).

### 3.5. Species delimitation, species validation and signals of introgression

Based on results from our population clustering and structure analyses, we tested up to 4 lineages (Western, Central, Eastern and Masirah) as new putative species selecting only non-admixed individuals. BFD* supported the species status for all four putative species with a decisive strength of support (ln(BF) > 5; Kass & Raftery, 1995; Table S5). Contrastingly, species delimitation analysis conducted with BPP supported the split of only three new putative species, lumping the Western and Central lineages into a single species (species delimitation model *H_2_* supported with a posterior probability of 0.95).

The heuristic genealogical divergence index (*gdi*) was implemented on *dataset 4 BPP* (Table S2), and similar results were recovered among both species tree topologies (Figure 4). First, *T. hajarensis* was compared to *T. spatalurus* to test the robustness of the current systematics on the group and results supported both species as distinct from each other with *gdi* values well above the 0.7 threshold (Figure 5c and Figure S5c). In contrast to the species delimitation models, comparisons between Hajar Mountain’s lineages did not support, in any case, the species status of these lineages, with *gdi* values falling either below 0.2 or within the ambiguous range. Masirah Island’s group presented a *gdi* above 0.9 which would support the species status of Masirah with respect to its sister group. Incongruently, Masirah’s sister lineage (either the Eastern Hajars lineage or the clade comprising all the Hajar Mountains’ *T. hajarensis*, depending on the topology) is not supported as a distinct species from Masirah (*gdi* = 0.19 0.01 in Figure 5c, and *gdi* = 0.35 0.09 in Figure S5c, respectively).

**Figure 5.**
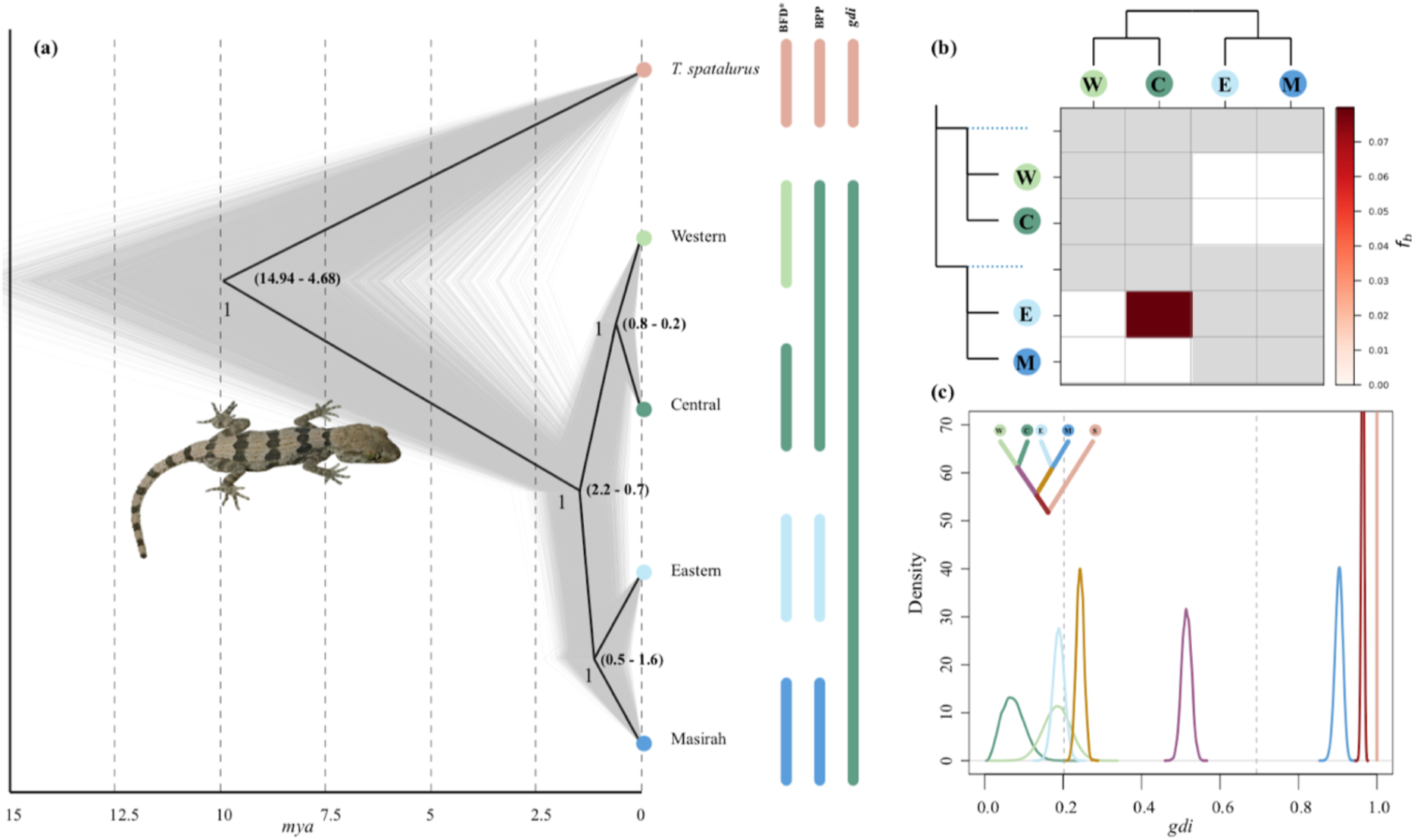
Species tree, gene flow and species delimitation analyses. **a:** Most supported species tree topology between SNAPP, BPP and IQ-TREE analyses. Species tree was inferred with SNAPP from *dataset 5 SNAPP* and both posterior (grey) and consensus trees (black) are shown. To the right, different coloured bars represent BFD*, BPP and *gdi* species level assignment; **b**: F-branch statistic analysis showing introgression between *Trachydactylus hajarensis* Eastern and Central clades; **c**: Posterior distribution for the *gdi* values between every pair of sister taxa within *Trachydactylus hajarensis*, including *T. spatalurus*. Colors in the internal branches of the upper-left corner species tree represent subsequent A00 BPP analyses where the descendant tips of the branch were lumped together and compared to its closest sister group. Values below 0.2 support a single species hypothesis while above 0.7 supports distinct species status.

Tests of introgression were implemented through D-statistics applied to the SNPs extracted from *dataset 3* (30,096 SNPs). When *Topo1* was given as a guide tree, statistically significant introgression was revealed between the Eastern and the Central lineages (Figure 5b). Otherwise, when *Topo2* was forced, significant past introgression was recovered between Eastern and Masirah Island lineages (Figure S5b).

## 4. Discussion

### 4.1. A highly structured species with past or recent gene flow

Our genomic analyses show that *T. hajarensis* is comprised by three well-defined lineages geographically located in the Western + Central Hajar Mountains, the Eastern Hajar Mountains, and in Masirah Island, respectively. However, the phylogenetic relationships of these lineages differ between methodologies showing high levels of mito-nuclear discordance (Figure 3) and introgression between either Central Hajars and Eastern Hajars lineages (Figure 5b), or Eastern Hajars and Masirah Island lineages (Figure S5b). The mitochondrial phylogenetic reconstruction showed a widely spread Central Hajars’ lineage including most of the Eastern Hajars, with only the easternmost region of the Hajar Mountains, near Ra’s al Hadd, being sister to the Masirah Island’s group (Figure 3). Incongruently, none of our nuclear genomic analyses recovered this topology, with a clear divergence between Central and Eastern lineages in the region surrounding the Semail gap (localities 19 to 21), a well-known topographic feature that has already been used to delimit the separation between the Eastern and Central Hajars (Garcia-Porta et al., 2017). Moreover, population structure analyses showed almost no signals of admixture between Central and Eastern populations (Figure 2a, b & c). However, when inspecting the fineRADstructure results (Figure 2e), the Eastern lineage appeared as greatly substructured, with an increasing gradient of shared alleles when geographically closer to the Central Hajars. This substructure may be caused by the habitat and climatic heterogeneity, and the complex topography of the Hajar Mountains (Burriel-Carranza et al., 2019; Carranza et al., 2018), which could be hindering panmixia within the group. The increasing gradient of coancestry observed in fineRADstructure also suggests a secondary contact between either the Eastern and Masirah lineages, or the Eastern and Central lineages, which was resolved by our MAST analysis (Figure 4). The MAST model recovered Masirah Island’s lineage as sister to the Eastern lineage and showed that the topology where all Hajar Mountains’ lineages form a monophyletic group (*Topo2*) is most likely a product of a secondary contact between Eastern and Central specimens, having a strong signal in the contact zone and gradually fading out with distance. The lack (or low levels) of current admixture between Eastern and Central specimens suggest that the observed patterns are a result of past secondary contacts or dispersal events from the Central Hajars towards the Eastern Hajars, leaving genomic evidence of nuclear and mitochondrial introgression between lineages.

Our D-statistic analysis points out that, among the number of processes leading to conflicts between mitochondrial and nuclear genomes, introgression might be responsible for the observed discordance in this case. Since the mitochondrial genome is maternally inherited and does not segregate or recombine, introgression events can incorporate a complete foreign mitochondrial genome into a population and maintain it over long periods of time, resulting into extremely diverged haplotypes between specimens from the same population but from differently-inherited mitochondrial genomes. Moreover, the lack (or low levels) of current nuclear admixture between Central and Eastern lineages support a past hybridization event.

### 4.2. Caveats of downsampling individuals when building species trees

The high computational resources needed to infer species trees with NGS data commonly leads to a dataset downsampling, selecting only some individuals for such analyses (Dufresnes et al., 2020; Kornilios et al., 2019; Thanou, Kornilios, Lymberakis, & Leaché, 2020). Moreover, since SNAPP does not assume gene flow, specimens are usually selected using population structure tools such as ADMIXTURE, with the assumption that any specimen from the same lineage species that is not admixed with other lineages should coalesce first within its group. Here, we provide an example where, even though discarding individuals with admixed proportions of their genomes, sample selection can greatly influence the topology and branch lengths of the resulting trees. It is especially worrying the case of the specimen OM04_2010_66-20, which even though it is not recovered as admixed in the best K of ADMIXTURE, it contains more loci supporting the hybrid topology (*Topo2*; Figure 4) than the *true* topology (*Topo1*; Figure 4). While most of the reconstructed species trees support *Topo1*, there are some cases where *Topo2* has higher node support (Figure S3).

In this case, several particular effects or the combination of them might be promoting such incongruent results. On the one hand, biological effects such as the small population size of Masirah Island, or the introgression between Eastern and Central+Western lineages might be confounding species tree estimation analyses. On the other hand, operational effects such as the usage of SNAPP linking all population sizes, a common feature used to reduce model parameters and achieve feasible run times (Stange et al., 2018) seems to be promoting a topology where Masirah Island is sister to the rest of the Hajar Mountains’ *T. hajarensis* (Figure S3a, c and d), while the same analyses unlinking population sizes recover a clearly distinct topology.

Overall, our study suggests that specimen selection when downsampling datasets for species tree estimation should be proceeded with caution, especially if there are signs of recent or past gene flow or different population sizes between taxa. Therefore, downsampled datasets should not include specimens near contact zones and, if possible, include specimens spanning throughout the whole distribution of each taxon.

### 4.3. One, two or three species? Importance of validation in species delimitation methods

Our analyses applying a variety of species delimitation methods yielded different results. Mitochondrial-based GMYC supports *T. hajarensis* to be a complex of five species, BFD* supports four species and BPP three. If we consider the integrated results of BFD* and BPP, both support the split of Western+Central, Eastern and Masirah as three distinct species. Such species would fall within the spectrum of cryptic species, since even though they are genetically identifiable there are no obvious distinctive phenotypic traits between them (de Pous et al. 2016; S.C. pers. observ.). Contrastingly, when using an heuristic criterion to validate species status (the *gdi*) none of the putative species was fully recovered as a distinct species (Figure 5c). The ambiguous results between Masirah Island’s lineage and its respective sister group are most likely due to a stated weakness of the *gdi* which, in the case of populations founded by a small number of individuals or in cases where two lineages have very different population sizes, results may lead to claims of species status even if the groups diverged very recently (Leaché et al., 2019). Our results support a single colonization event of Masirah Island and the high number of shared alleles between specimens also suggests that this population was founded by few individuals. Considering the different population sizes with its sister group and the low values of *gdi* of its counterpart test (Figure 5c), we identify Masirah Island’s specific status claim as a false positive. Validating SDMs has proven crucial to not contribute to taxonomic inflation in *T. hajarensis* and we suggest that, together with an in-depth exploration of population structure and gene flow, the *gdi* (or any other similar heuristic method) should become a key tool in species delimitation validation, specially when testing the species status of allopatric early divergent lineages (Leaché et al., 2019).

### 4.4. Systematics and Biogeography

The present study shows that *T. hajarensis* is formed by three well supported lineages with a complex and dynamic evolutionary history. The first divergence within the species separated the *T. hajarensis* currently inhabiting the Western and Central Hajar Mountains from the *T. hajarensis* on Masirah Island and the Eastern Hajars. The split between these clades is estimated around Early to Mid-Quaternary (1.4 my; 0.7-2.13 mya HPD95%; Figure 5), setting the first diversification within the group about three million years after previous estimations using mitochondrial markers (de Pous et al., 2016; Figure 3). Given the distribution of the different lineages we can assume that they speciated through allopatric isolation, probably caused by a combination of past geographical and climatic events. Interestingly, Western + Central Hajars lineage and Eastern hajars lineage are found nowadays in allopatry in the same mountain range (Figure 1). However, the introgression signal recovered between the two lineages and the mitochondrial phylogeny suggest that at some point they cohabited and gene flow between lineages occurred. This could indicate that the Eastern lineage colonized the Hajar Mountains posteriorly to the Western + Central clade and progressively displaced the Western + Central populations to its current distribution exchanging genes in the process. Another explanation would be that Western + Central populations in the Eastern Hajars merged with the arriving Eastern lineage and the high topography of the Central Hajars hindered the dispersal and complete fusion of both lineages into one. There are other examples where genetic isolation is found between lineages in the Eastern and Central Hajars (Carranza & Arnold, 2012; Garcia-Porta et al., 2017) and in the case of the geckos of the genus *Asaccus*, it has even led to the description of a new species (Simó-Riudalbas et al., 2018). Overall, we can speculate that even though currently there is low gene flow between lineages, genetic barriers are diffuse and, if at some point climatic conditions favor increased dispersal of this ground-dwelling species, both groups could eventually fuse back into one.

In the Arabian Peninsula, Pliocene and Quaternary climatic shifts between humid and arid to hyper-arid episodes have resulted from repeated high-latitude glacial events and global sea level falls, promoting desert formation (Glennie, 1998). In southeastern Oman, we find the Sharqiyah Sands, a sand dune desert where *T. hajarens*is has never been reported and which separates the Masirah’s lineage by roughly 200 km in a straight line to its closest sister group in the Eastern Hajars (Figure 1). This desert is thought to be a result of Late Quaternary climatic shifts in southeast Arabia (although see Metallinou & Carranza, 2013) and its formation is linked to the onshore-blowing SW Monsoon (Glennie, 1998) with prior fluvial deposits dating back to the Plio-Pleistocene and first aeolian sands consolidating about 160,000 years ago (Radies, Preusser, Matter, & Mange, 2004). The divergence between Masirah and Eastern lineages dates to 1.1 mya (0.5 – 1.6 mya HPD95%; Figure 5), a time range prior to the consolidation of the Sharqyah Sands. This would be coherent with a broader historic distribution range of *T. hajarensis* encompassing the current region where today the Sharqyah Sands reside. This would have facilitated colonization by adrift specimens or clutches, since Masirah Island is only about 20 km from the closest continental shore. Moreover, sea level drops of up to 130 m induced by high-latitude glacial events were recurrent during the Plio-Pleistocene (Gleenie, 1998). The greatest depth between Masirah Island and the coast of Oman rarely surpasses 50 m, thus such sea level drops could have provided a land bridge between Masirah Island and the continent, facilitating land dispersal of *T. hajarensis* and other reptile species towards the Island. Phylogenomic reconstructions and population structure analyses support a single colonization event of Masirah Island from few founder individuals, with all specimens constituting a monophyletic group (Figure 3b) with high levels of coancestry among them (Figure 2d). These results, together with the increased sampling effort throughout several localities in the island, are in agreement with previous findings (de Pous et al., 2016), and also advocate for a natural colonization of Masirah Island in contrast to a human mediated introduction.

## 5. Conclusions

Overall, *T. hajarensis* seems to be conformed by a single, highly-structured species. The usage of genomic techniques has proven vital to determine the population structure, introgression events, and mito-nuclear discordances within this system, but further studies with whole genome sequencing data might be necessary to fully understand the regions of introgression within the species. Moreover, although NGS and the MSC are essential tools to shorten the Linnean Shortfall and are well suited to uncover cryptic diversity, we show that dataset downsampling and species delimitation methods need to be cautiously implemented and validated to avoid contributing to taxonomic inflation. Altogether, this system offers the rare possibility to witness both possible evolutionary outcomes of an incipient species at the same time. On the one hand, if isolation is maintained between continental and Masirah Island’s populations, both lineages will most likely evolve into different species. On the other hand, its sister lineage could potentially merge back with the other Hajar Mountain’s *T. hajarensis*, dissolving almost 1.5 my of independent evolution. However, only time will tell.

## Author’s contributions

BB-C and SC conceived and designed the study. BB-C and ME performed laboratory work, analyzed the data and wrote the manuscript with input from all other authors, who approved the paper in its final form.

## Aknowledgements

We would like to thank all the past and present members of the Ministry of Environment and Climate Affairs, MECA, Oman, now the Environment Authority Oman, and especially to Ali Al Kiyumi, Suleiman Nasser Al Akhzami, Thuraya Al Sariri, Ahmed Said Al Shukaili, Ali Alghafri, Sultan Khalifa, Hamed Al Farqani, Salim Bait Bilal, Iman Sulaiman Alzari, Aziza Saud Al Adhoobi, Mohammed Al Shariani, Zeyana Salim Al Omairi and Abdullah bin Ali Al Amri, Chariman of the Environment Authority. We are also very grateful to past members and collaborators including Edwin Nicholas Arnold, Michael D. Robinson, Andrew Gardner, Josep Roca, Philip de Pous, Margarita Metallinou, Loukia Spilani and Dean Addams for their help and support. In the UAE, we wish to thank His Highness Sheikh Dr. Sultan bin Mohammed Al Qasimi, Supreme Council Member and Ruler of Sharjah, H. E. Ms. Hana Saif al Suwaidi (Chairperson of the Environment and Protected Areas Authority, Sharjah), Paul Vercammen and Kevin Budd (Breeding Centre for Endangered Arabian Wildlife), and Gary Feulner (Dubai Natural History Group) for their continuous support. SC is supported by grants PGC2018-098290-B-I00 (MCIU/AEI/FEDER, UE) and PID2021-128901NB-I00 funded by MCIN/AEI/ 10.13039/501100011033 and by ERDF, a way of making Europe. BB-C was funded by FPU grant from Ministerio de Ciencia, Innovación y Universidades, Spain (FPU18/04742). AT is supported by “la Caixa” doctoral fellowship programme (LCF/BQ/DR20/11790007). GR was funded by an FPI grant from the Ministerio de Ciencia, Innovación y Universidades, Spain (PRE2019-088729). HT-C is supported by a “Juan de la Cierva - Formación” postdoctoral fellowship (FJC2021-046832-I) funded by MCIN/AEI/10.13039/501100011033 and by the European Union NextGenerationEU/PRTR. No in vivo experiments were performed. The field study was carried out with the authorization of the governments of UAE and Oman. Permits from Oman were issued by the Nature Conservation Department of the Ministry of Environment and Climate Affairs, Oman (Refs: 08/2005; 16/2008; 38/2010; 12/2011; 13/2013; 21/2013; 37/2014; 31/2016; 6210/10/21).

## SUPPLEMENTARY FIGURES

**Figure S1.**
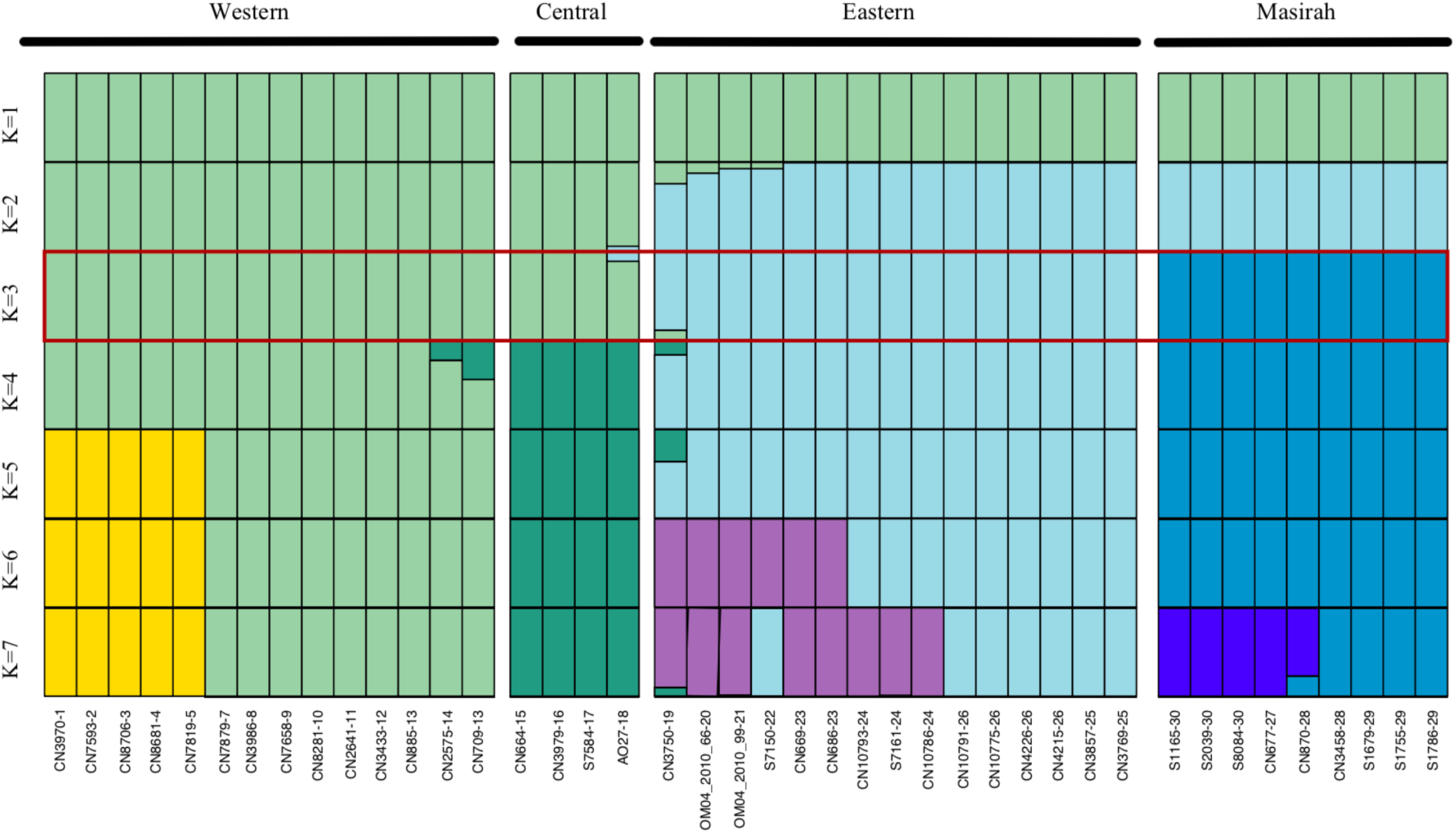
Individual assignments of ADMIXTURE nuclear clusters from K = 1 to K = 7. The lowest cross validation result, depicting the most probable number of populations, is highlighted in red. Numbers after last dash correspond to locality number from Figure 1 and Table S1. Information on all the specimens analyzed can be found in Table S1.

**Figure S2.**
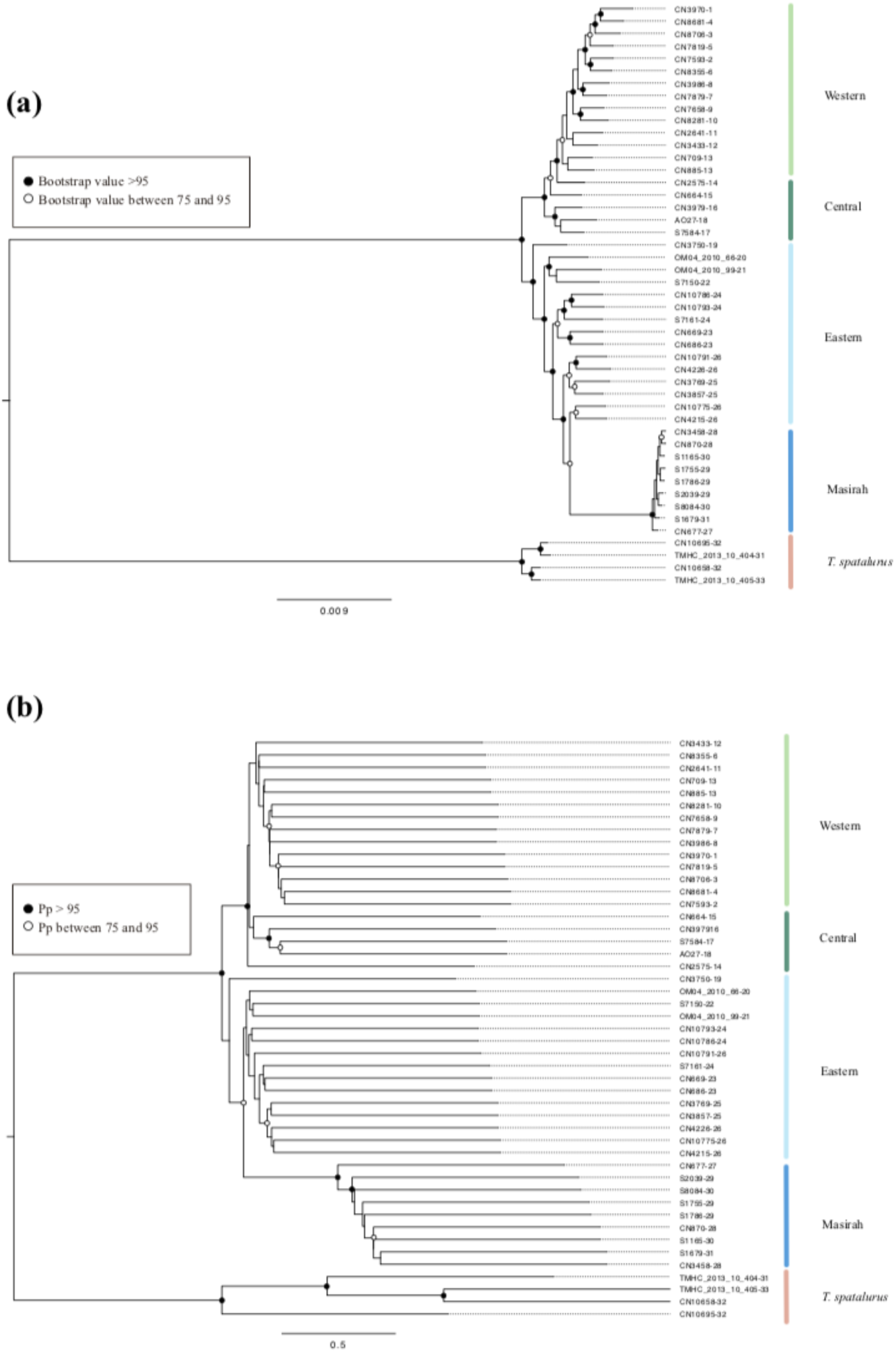
Maximum likelihood nuclear phylogenomic reconstructions. **a**: Tree inferred with RAxML-NG with a concatenated dataset of 5,219 loci; **b**: Consensus tree inferred with Astral after computing 5,219 gene trees in IQ-TREE. Western, Central, Eastern and Masirah represent the different *Trachydactylus hajarensis* clades. Numbers after last dash correspond to locality number from Figure 1 and Table S1.

**Figure S3.**
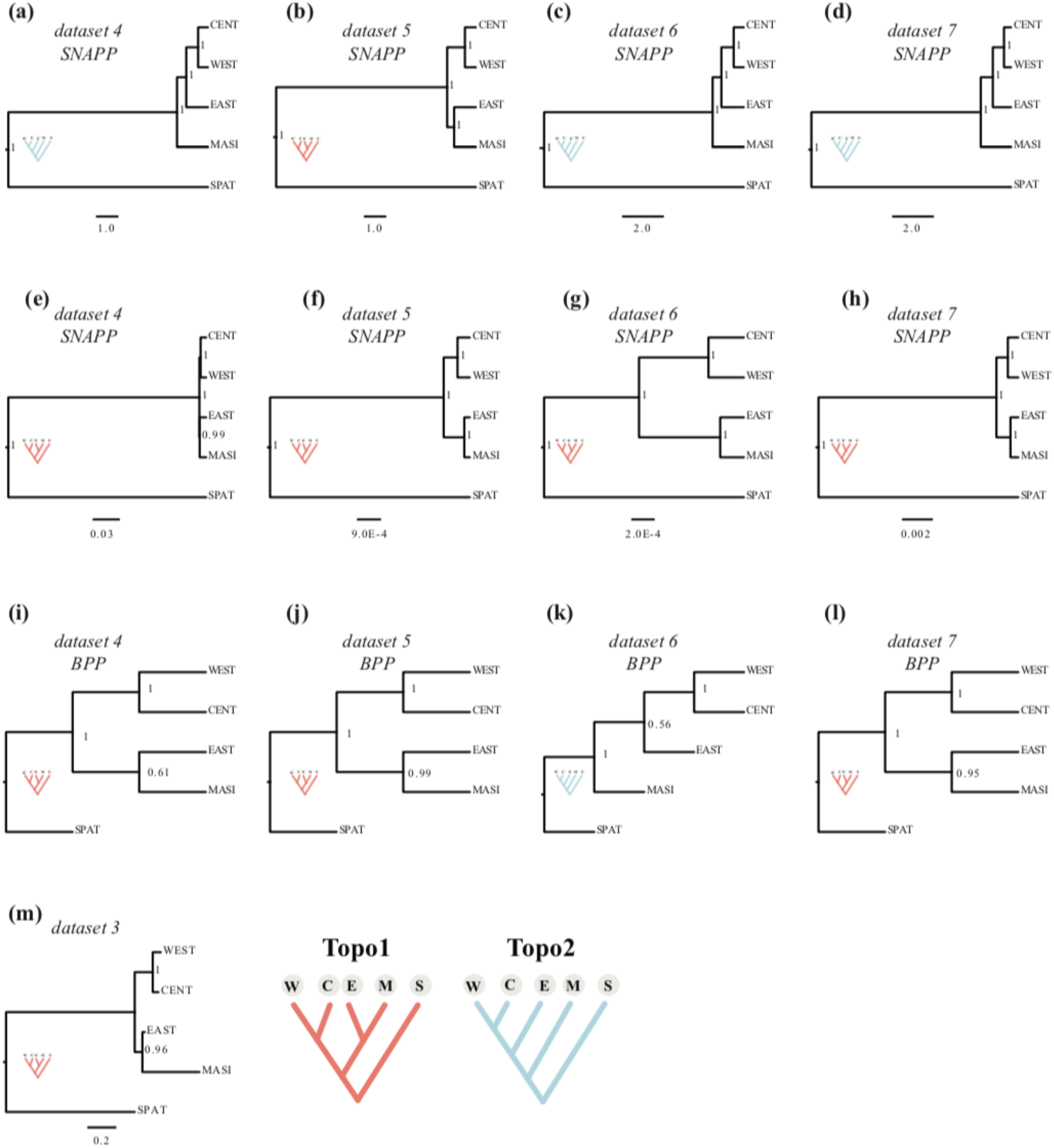
Species tree topologies with different sample selection. Samples included in each analysis range between 10–20 specimens in *datasets 4–7*, and 47 samples (all available samples) in *dataset 3*. For more information on each dataset see Table S2. Species trees were inferred with SNAPP linking all population sizes (**a**–**d**), with SNAPP without linking population sizes (**e**–**h**), with BPP A01 approach (**i**–**l**), and with Astral after summarizing all 5,219 gene trees obtained with IQtree (**m**). Abbreviations are as follow: WEST, Western Clade of *T. hajarensis*; CENT, Central Clade of *T. hajarensis*; EAST, Eastern Clade of *T. hajarensis*; MASI, Masirah Island Clade of *T. hajarensis;* SPAT, *T. spatalurus*.

**Figure S4.**
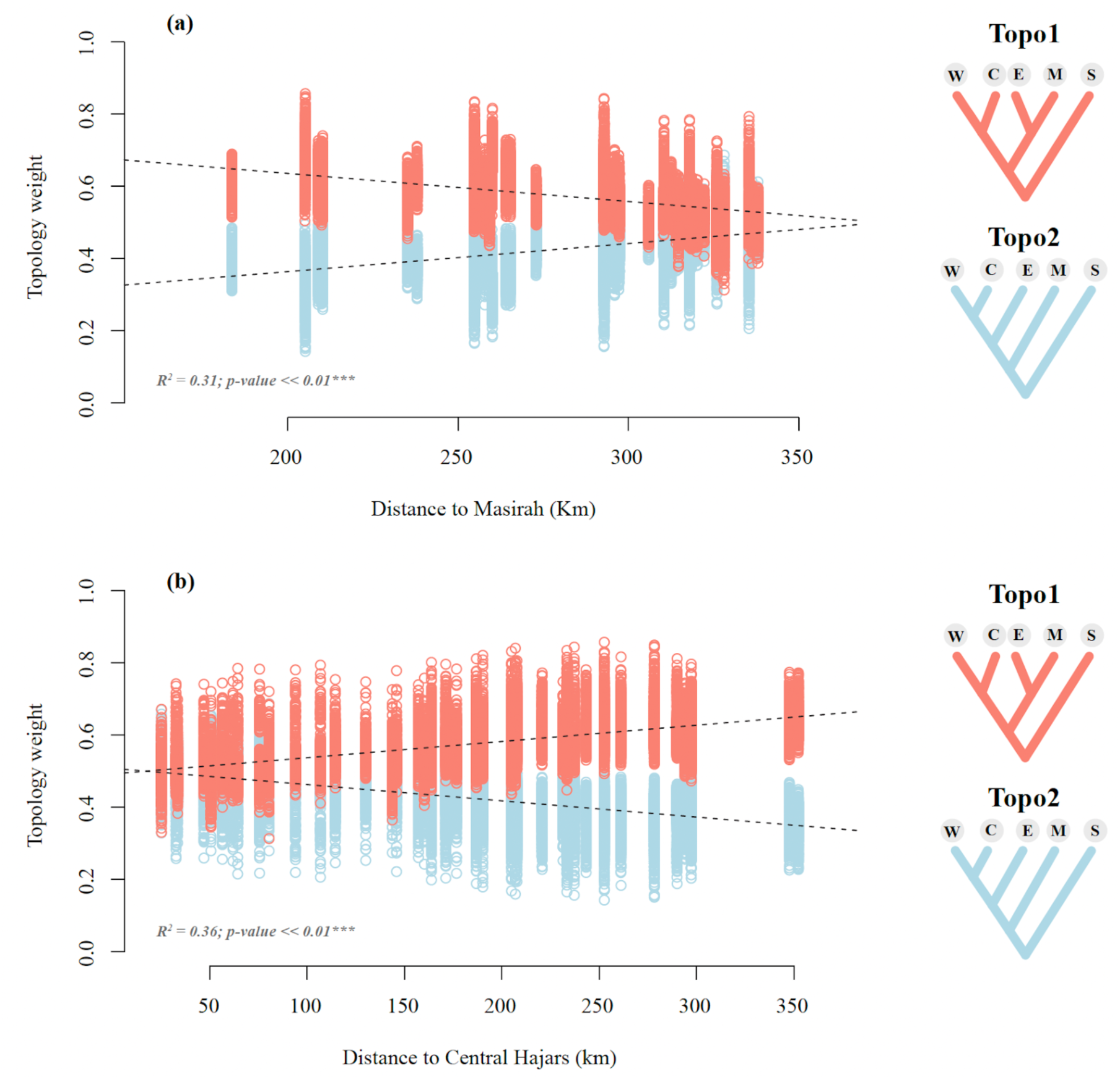
**a:** Shifts in *Topo1* (salmon) and *Topo2* (blue) topology weights explained by the distance from each Eastern Hajars’ specimen to each Masirah Island’s specimen; **b:** Shifts in *Topo1* (salmon) and *Topo2* (blue) topology weights explained by the distance from each Eastern Hajars’ specimen to each Central Hajars’ individual. Each point represents the averaged topology weight of a quintet across all its loci. Points distributed across vertical lines correspond to quintet combinations where Eastern and Masirah Island specimens remained identical but the individuals from the other lineages varied. We accounted for such pseudoreplicates by averaging each locality’s topology weight (Figure 3), or by including them into a nested ANOVA (Tables S3 and S4). The sum of topology weights (*Topo1* and *Topo2*) of each MAST analysis always adds up to 1.0, thus plotted regression lines have inverse directions but identical R^2^ and p-values.

**Figure S5.**
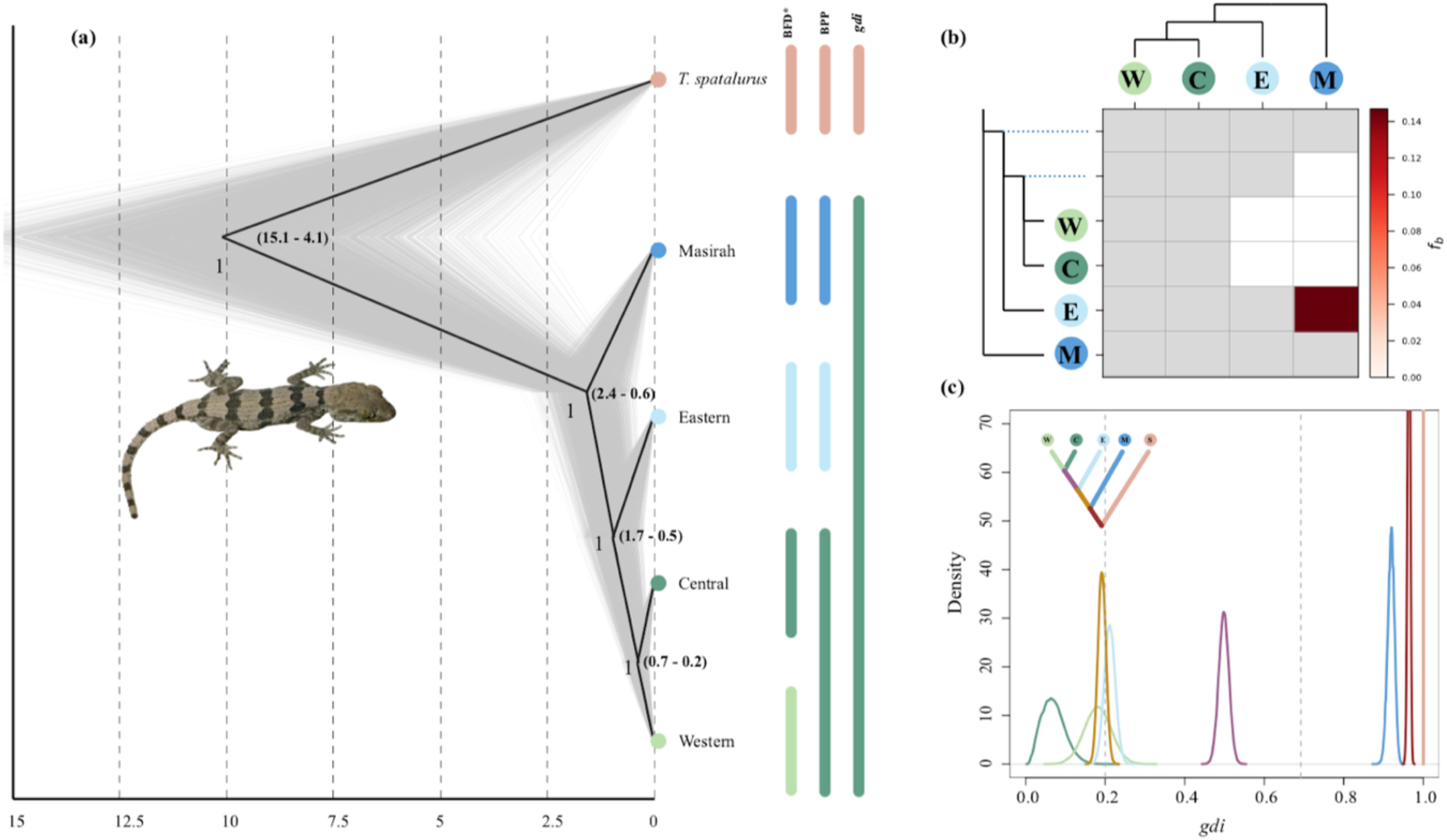
Species tree, gene flow and species delimitation analyses. **a:** Second species tree topology obtained with SNAPP & BPP. Species tree was inferred with SNAPP from *dataset* 4 *SNAPP* and posterior trees (grey) and consensus tree (black) are shown. To the right, different coloured bars represent BFD*, BPP and *gdi* species level assignment, respectively; **b**: F-branch statistic analysis showing introgression between *Trachydactylus hajarensis* Eastern and Masirah Island clades; **c**: Posterior distribution for the *gdi* values between every pair of sister taxa within *Trachydactylus hajarensis*, including *T. spatalurus*. Colors in the internal branches of the upper-left corner species tree represent subsequent A00 BPP analyses where the descendant tips of the branch were lumped together and compared to its closest sister group. Values below 0.2 support a single species hypothesis while above 0.7 supports distinct species status.

## SUPPLEMENTARY TABLES

**Table S1.**
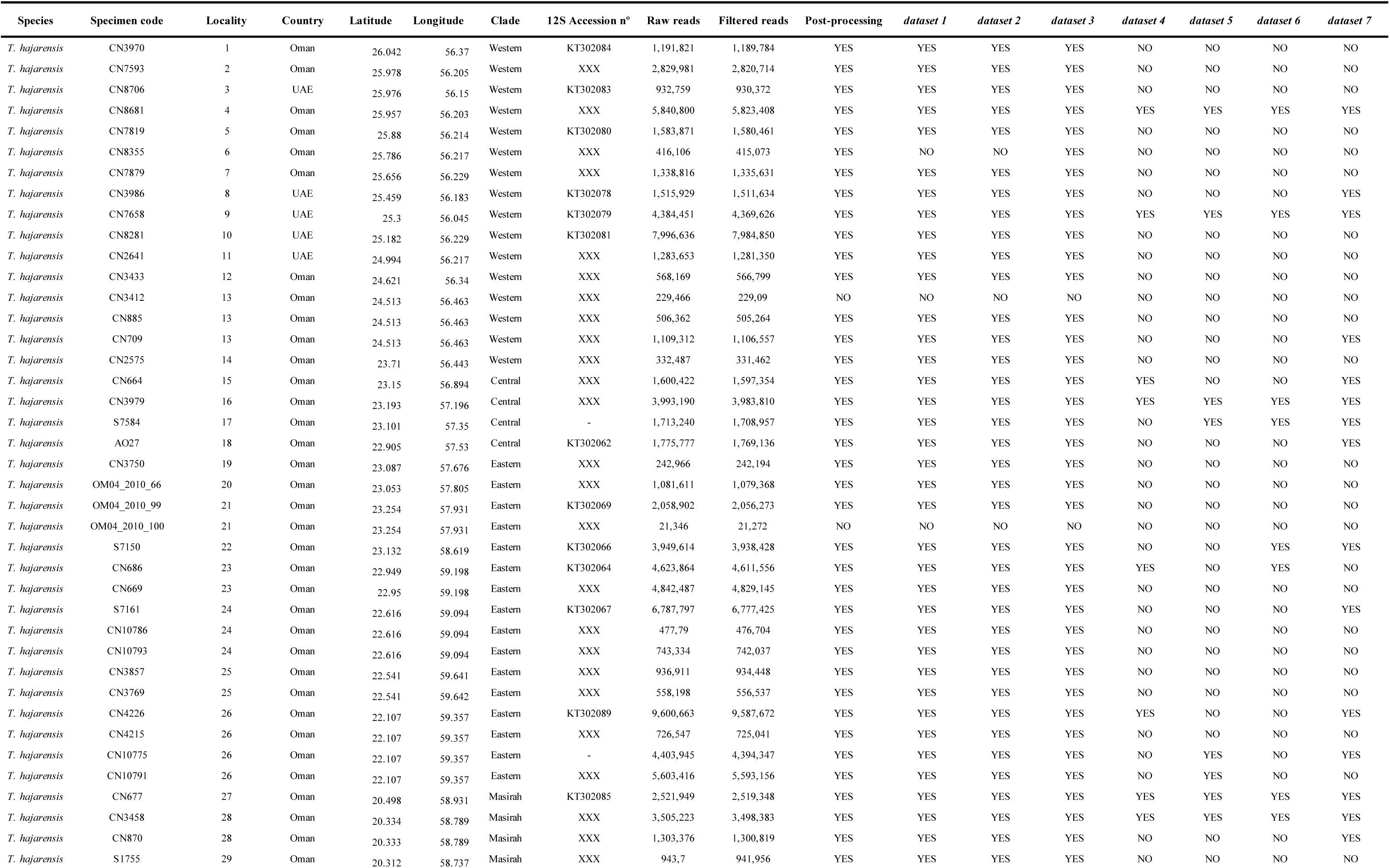
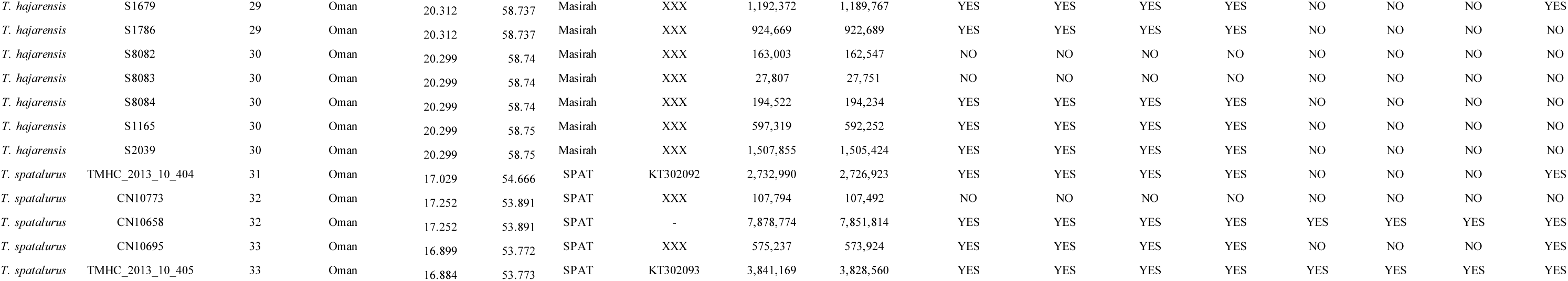
Table of all specimens included in this study information regarding location, clade, GenBank accession number, ddRADseq raw and filtered reads as well as information on which individuals are included in each dataset. GenBank accession numbers labled XXXX were de novo produced and will be uploaded; datasets 4-7 include both SNAPP and BPP datasets in Table S2.

**Table S2:** dataset specifications. uSNPs: Unlinked Single Nucleotide Polymorphisms; Ind: Missing data allowed per individual; gen: Missing genotype call rate allowed; bp: Base pairs.

| dataset name | dataset type | Analysis | Postprocessing filtering (%) | n° of individuals | dataset length (bp) | n° of loci | n° of SNPs | Missingness (%) |
| --- | --- | --- | --- | --- | --- | --- | --- | --- |
| *dataset 1* | *uSNPs* | AdMIXTURE & PCA | 78 ind – 40 gen | 42 | 2,428 | 2,428 | 2,428 | 18.57 |
| *dataset 2* | *loci* | fineRADstructure | Not filtered | 42 | 1,973,898 | 30,526 | 99,828 | 63.98 |
| *dataset 3* | *loci* | raxml-ng, IQTREE, BEAST, SplitsTree, MAST & dsuite | 78 ind – 40 gen | 47 | 338,507 | 5,219 | 30,096 | 22.21 |
| *dataset 4 SNAPP* | *uSNPs* | Species tree inference and Species delimitation | 10 gen | 10 | 2,147 | 2,147 | 2,147 | 4.23 |
| *dataset 5 SNAPP* | *uSNPs* | Species tree inference | 10 gen | 10 | 3,210 | 3,210 | 3,210 | 4.88 |
| *dataset 6 SNAPP* | *uSNPs* | Species tree inference | 10 gen | 10 | 2,897 | 2,897 | 2,897 | 4.84 |
| *dataset 7 SNAPP* | *uSNPs* | Species tree inference | 10 gen | 20 | 2,731 | 2,731 | 2,731 | 9.72 |
| *dataset 4 BPP* | *loci* | Species tree inference, Species delimitation and *gdi* | 60 gen | 10 | 261,420 | 4,357 | 20,246 | 15.54 |
| *dataset 5 BPP* | *loci* | Species tree inference | 60 gen | 10 | 271,980 | 4,533 | 19,966 | 13.7 |
| *dataset 6 BPP* | *loci* | Species tree inference | 60 gen | 10 | 266,460 | 4,441 | 20,262 | 15.3 |
| *dataset 7 BPP* | *loci* | Species tree inference | 60 gen | 20 | 315,240 | 5,254 | 26,103 | 27.2 |

**Table S4.** Nested analysis of variance table. Response variable shown in this nested ANOVA correspond to topology weigths 1. However, the exact same results were obtanied when using topology weigths 2. Dist ME: Distance between Masirah and Eastern specimens

| **Response Topo1** | **DF** | **Sum_sq** | **Mean_Sq** | **F value** | **Pr(>F)** |
| --- | --- | --- | --- | --- | --- |
| *Dist ME* | 1 | 45.892 | 45.89 | 11,399.25 | << 0.01 |
| *Dist ME:replicate* | 3,359 | 13.523 | 0.004 | 1,585 | << 0.01 |
| *Residuals* | 34,439 | 87.472 | 0.003 |  |  |

**Table S4.** Nested analysis of variance table. Response variable shown in this nested ANOVA correspond to topology weigths 1. However, the exact same results were obtanied when using topology weigths 2. Dist CE: Distance between Central and Eastern specimens

| **Response Topo1** | **DF** | **Sum_sq** | **Mean_Sq** | **F value** | **Pr(>F)** |
| --- | --- | --- | --- | --- | --- |
| *Dist CE* | 1 | 53.144 | 53.144 | 3,801.42 | << 0.01 |
| *Dist CE:replicate* | 2,015 | 28.17 | 0.014 | 7.629 | << 0.01 |
| *Residuals* | 35,783 | 65.572 | 0.002 |  |  |

**Table S5:**
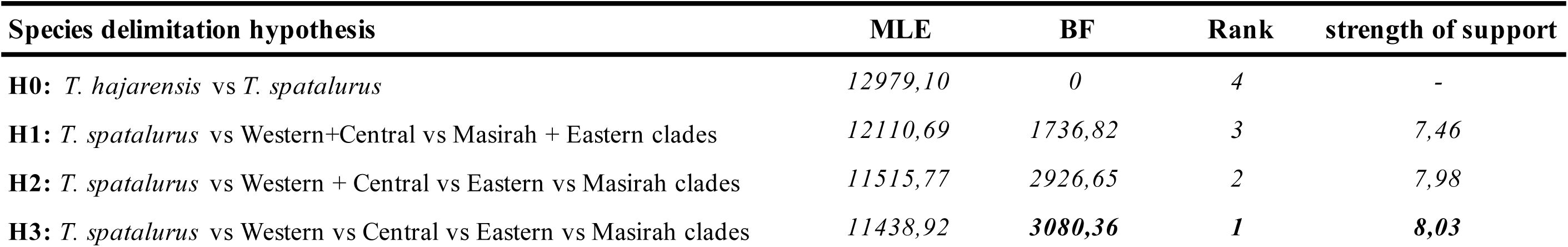
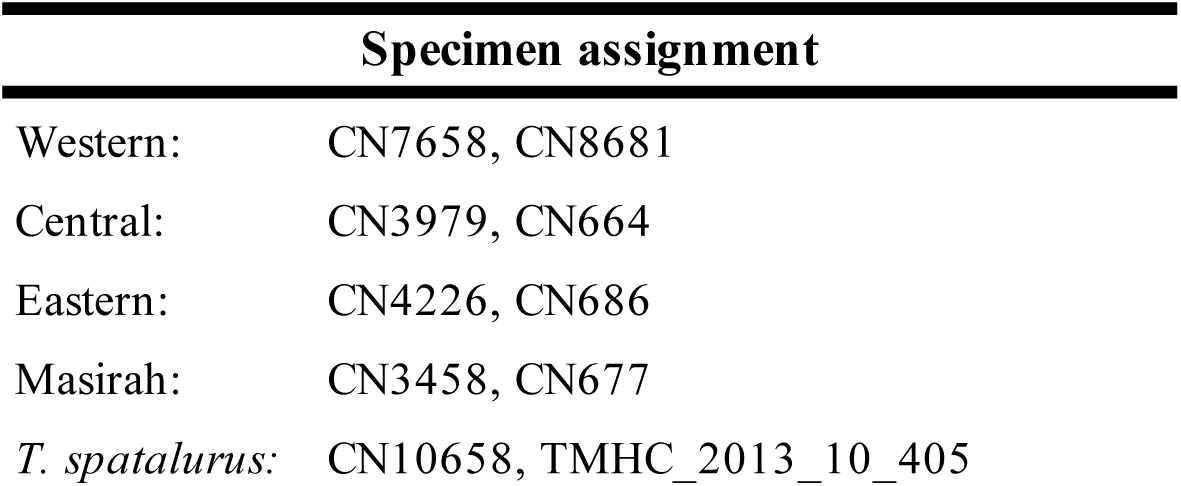
Results of BFD* testing the support of the competing species hypotheses. For each hypothesis we show Maximum Likelihood Estimates (MLE), their Bayes Factors (*2 x (H0 - Hi)*), the rank and the strength of support *ln(BF).* Additionally, we show the specimens assigned to each putative species hypothesis.

## Notes

### Competing Interest Statement

The authors have declared no competing interest.

## References

Arnold, E. N. (1980). The reptiles and amphibians of Dhofar, southern Arabia. Journal of Oman Studies, Special Report, 2, 273–332.

Bouckaert, R., Vaughan, T. G., Barido-Sottani, J., Duchêne, S., Fourment, M., Gavryushkina, A., … Drummond, A. J. (2019). BEAST 2.5: An advanced software platform for Bayesian evolutionary analysis. PLOS Computational Biology, 15(4), e1006650. doi: 10.1371/JOURNAL.PCBI.1006650

Bray, T. C., Alagaili, A. N., & Bennett, N. C. (2014). A widespread problem: Cryptic diversity in the Libyan jird. Zoological Studies 2014 53:1, 53(1), 1–8. doi: 10.1186/S40555-014-0033-3

Bryant, D., Bouckaert, R., Felsenstein, J., Rosenberg, N. A., & Roychoudhury, A. (2012). Inferring Species Trees Directly from Biallelic Genetic Markers: Bypassing Gene Trees in a Full Coalescent Analysis. Molecular Biology and Evolution, 29(8), 1917–1932. doi: 10.1093/MOLBEV/MSS086

Burriel-Carranza, B., Els, J., & Carranza, S. (2022). Reptiles & Amphibians of the Hajar Mountains. Madrid: Editorial CSIC. ISBN: 978-84-00-10988-2.

Burriel-Carranza, B., Tarroso, P., Els, J., Gardner, A., Soorae, P., Mohammed, A. A., … Carranza, S. (2019). An integrative assessment of the diversity, phylogeny, distribution, and conservation of the terrestrial reptiles (Sauropsida, Squamata) of the United Arab Emirates. PLOS ONE, 14(5), e0216273. doi: 10.1371/JOURNAL.PONE.0216273

Carranza, S., & Arnold, E. N. (2012). A review of the geckos of the genus *Hemidactylus* (Squamata: Gekkonidae) from Oman based on morphology, mitochondrial and nuclear data, with descriptions of eight new species. In Zootaxa (Vol. 95).

Carranza, S., Els, J., & Burriel-Carranza, B. (2021). A field guide to the reptiles of Oman. Madrid: Digital CSIC. ISBN: 978-84-00-10876-2.

Carranza, S., Xipell, M., Tarroso, P., Gardner, A., Arnold, E. N., Robinson, M. D., … Akhzami, S. N. A. (2018). Diversity, distribution and conservation of the terrestrial reptiles of Oman (Sauropsida, Squamata). In PLoS ONE (Vol. 13). doi: 10.1371/journal.pone.0190389

Chan, K. O., Hutter, C. R., Wood, P. L., Grismer, L. L., Das, I., & Brown, R. M. (2020). Gene flow creates a mirage of cryptic species in a Southeast Asian spotted stream frog complex. Molecular Ecology, 29(20), 3970–3987. doi: 10.1111/mec.15603

Chang, C. C., Chow, C. C., Tellier, L. C. A. M., Vattikuti, S., Purcell, S. M., & Lee, J. J. (2015). Second-generation PLINK: Rising to the challenge of larger and richer datasets. GigaScience, 4(1), 7. doi: 10.1186/S13742-015-0047-8/2707533

Chattopadhyay, B., Garg, K. M., Kumar, A. K. V., Doss, D. P. S., Rheindt, F. E., Kandula, S., & Ramakrishnan, U. (2016). Genome-wide data reveal cryptic diversity and genetic introgression in an Oriental cynopterine fruit bat radiation. BMC Evolutionary Biology, 16(1), 1–15. doi: 10.1186/S12862-016-0599-Y/TABLES/1

de Pous, P., Machado, L., Metallinou, M., Červenka, J., Kratochvíl, L., Paschou, N., … Carranza, S. (2016). Taxonomy and biogeography of *Bunopus spatalurus* (Reptilia; Gekkonidae) from the Arabian Peninsula. Journal of Zoological Systematics and Evolutionary Research, 54(1), 67–81. doi: 10.1111/jzs.12107

De Queiroz, K. (2007). Species Concepts and Species Delimitation. Systematic Biology, 56(6), 879–886. doi: 10.1080/10635150701701083

Derycke, S., Fonseca, G., Vierstraete, A., Vanfleteren, J., Vincx, M., & Moens, T. (2008). Disentangling taxonomy within the *Rhabditis* (*Pellioditis*) *marina* (Nematoda, Rhabditidae) species complex using molecular and morhological tools. Zoological Journal of the Linnean Society, 152(1), 1–15. doi: 10.1111/J.1096-3642.2007.00365.X

Dufresnes, C., Pribille, M., Alard, B., Gonçalves, H., Amat, F., Crochet, P.-A., … Martínez-Solano, I. (2020). Integrating hybrid zone analyses in species delimitation: Lessons from two anuran radiations of the Western Mediterranean. Heredity, 124(3), 423–438. doi: 10.1038/s41437-020-0294-z

Eaton, D. A. R., & Overcast, I. (2020). ipyrad: Interactive assembly and analysis of RADseq datasets. Bioinformatics, 36(8), 2592–2594. doi: 10.1093/BIOINFORMATICS/BTZ966

Ezard, T., Fujisawa, T., & Barraclough, T. (2021). splits: SPecies’ LImits by Threshold Statistics. Retrieved from https://R-Forge.R-project.org/projects/splits/

Fennessy, J., Bidon, T., Reuss, F., Kumar, V., Elkan, P., Nilsson, M. A., … Janke, A. (2016). Multi-locus Analyses Reveal Four Giraffe Species Instead of One. Current Biology, 26(18), 2543–2549. doi: 10.1016/J.CUB.2016.07.036

Fišer, C., Robinson, C. T., & Malard, F. (2018). Cryptic species as a window into the paradigm shift of the species concept. Molecular Ecology, 27(3), 613–635. doi: 10.1111/MEC.14486

Flouri, T., Jiao, X., Rannala, B., & Yang, Z. (2018). Species Tree Inference with BPP Using Genomic Sequences and the Multispecies Coalescent. Molecular Biology and Evolution, 35(10), 2585–2593. doi: 10.1093/molbev/msy147

Fujisawa, T., & Barraclough, T. G. (2013). Delimiting Species Using Single-Locus Data and the Generalized Mixed Yule Coalescent Approach: A Revised Method and Evaluation on Simulated Data Sets. Systematic Biology, 62(5), 707–724. doi: 10.1093/sysbio/syt033

Garcia-Porta, J., Simó-Riudalbas, M., Robinson, M., & Carranza, S. (2017). Diversification in arid mountains: Biogeography and cryptic diversity of *Pristurus rupestris rupestris* in Arabia. Journal of Biogeography, 44(8), 1694–1704. doi: 10.1111/jbi.12929

Glennie, K. W. (1998). The desert of southeast Arabia: A product of Quaternary climatic change. In Quaternary Deserts and Climatic Change. CRC Press.

Gosselin, T., Lamothe, M., Devloo-Delva, F., & Grewe, P. (2017). radiator: RADseq data exploration, manipulation and visualization using R. R Package Version 0.0, 5.

Huson, D. H., & Bryant, D. (2006). Application of Phylogenetic Networks in Evolutionary Studies. Molecular Biology and Evolution, 23(2), 254–267. doi: 10.1093/molbev/msj030

Isaac, N. J. B., Mallet, J., & Mace, G. M. (2004). Taxonomic inflation: Its influence on macroecology and conservation. Trends in Ecology & Evolution, 19(9), 464–469. doi: 10.1016/J.TREE.2004.06.004

Ivanov, V., Lee, K. M., & Mutanen, M. (2018). Mitonuclear discordance in wolf spiders: Genomic evidence for species integrity and introgression. Molecular Ecology, 27(7), 1681–1695. doi: 10.1111/MEC.14564

Jackson, N. D., Morales, A. E., Carstens, B. C., & O’Meara, B. C. (2017). PHRAPL: Phylogeographic Inference Using Approximate Likelihoods. Systematic Biology, 66(6), 1045–1053. doi: 10.1093/SYSBIO/SYX001

Kass, R. E., & Raftery, A. E. (1995). Bayes Factors. Journal of the American Statistical Association, 90(430), 773–795. doi: 10.1080/01621459.1995.10476572

Knaus, B. J., & Grünwald, N. J. (2017). vcfr: A package to manipulate and visualize variant call format data in R. Molecular Ecology Resources, 17(1), 44–53. doi: 10.1111/1755-0998.12549

Kornilios, P., Thanou, E., Lymberakis, P., Ilgaz, Ç., Kumlutaş, Y., & Leaché, A. (2019). Genome-wide markers untangle the green-lizard radiation in the Aegean Sea and support a rare biogeographical pattern. Journal of Biogeography, 46(3), 552–567. doi: 10.1111/jbi.13524

Kozlov, A. M., Darriba, D., Flouri, T., Morel, B., & Stamatakis, A. (2019). RAxML-NG: a fast, scalable and user-friendly tool for maximum likelihood phylogenetic inference. Bioinformatics, 35(21), 4453–4455. doi: 10.1093/BIOINFORMATICS/BTZ305

Leaché, A. D., Zhu, T., Rannala, B., & Yang, Z. (2019). The Spectre of Too Many Species. Systematic Biology, 68(1), 168–181. doi: 10.1093/SYSBIO/SYY051

Main, D. C., van Vuuren, B. J., Tilbury, C. R., & Tolley, K. A. (2022). Out of southern Africa: Origins and cryptic speciation in *Chamaeleo*, the most widespread chameleon genus. Molecular Phylogenetics and Evolution, 175, 107578. doi: 10.1016/J.YMPEV.2022.107578

Malinsky, M., Matschiner, M., & Svardal, H. (2021). Dsuite—Fast D-statistics and related admixture evidence from VCF files. Molecular Ecology Resources, 21(2), 584–595. doi: 10.1111/1755-0998.13265

Marshall, T. L., Chambers, E. A., Matz, M. V., & Hillis, D. M. (2021). How mitonuclear discordance and geographic variation have confounded species boundaries in a widely studied snake. Molecular Phylogenetics and Evolution, 162, 107194. doi: 10.1016/J.YMPEV.2021.107194

Mayden, R. L. (1997). A hierarchy of species concepts: The denouement in the saga of the species problem (Claridge M.F, Dawah H.A, & Wilson M.R., Eds.). Retrieved from https://philpapers.org/rec/MAYAHO-6

Metallinou, M., & Carranza, S. (2013). New species of *Stenodactylus* (Squamata: Gekkonidae) from the Sharqiyah Sands in northeastern Oman. Zootaxa, 3745, 449–468. doi: 10.11646/zootaxa.3745.4.3

Metallinou, M., Červenka, J., Crochet, P. A., Kratochvíl, L., Wilms, T., Geniez, P., … Carranza, S. (2015). Species on the rocks: Systematics and biogeography of the rock-dwelling *Ptyodactylus geckos* (Squamata: Phyllodactylidae) in North Africa and Arabia. Molecular Phylogenetics and Evolution, 85, 208–220. doi: 10.1016/j.ympev.2015.02.010

Nguyen, L.-T., Schmidt, H. A., von Haeseler, A., & Minh, B. Q. (2015). IQ-TREE: A Fast and Effective Stochastic Algorithm for Estimating Maximum-Likelihood Phylogenies. Molecular Biology and Evolution, 32(1), 268–274. doi: 10.1093/molbev/msu300

O’Leary, S. J., Puritz, J. B., Willis, S. C., Hollenbeck, C. M., & Portnoy, D. S. (2018). These aren’t the loci you’e looking for: Principles of effective SNP filtering for molecular ecologists. Molecular Ecology, 27(16), 3193–3206. doi: 10.1111/MEC.14792

Peterson, B. K., Weber, J. N., Kay, E. H., Fisher, H. S., & Hoekstra, H. E. (2012). Double Digest RADseq: An Inexpensive Method for De Novo SNP Discovery and Genotyping in Model and Non-Model Species. PLOS ONE, 7(5), e37135. doi: 10.1371/JOURNAL.PONE.0037135

Pons, J., Barraclough, T. G., Gomez-Zurita, J., Cardoso, A., Duran, D. P., Hazell, S., … Vogler, A. P. (2006). Sequence-Based Species Delimitation for the DNA Taxonomy of Undescribed Insects. Systematic Biology, 55(4), 595–609. doi: 10.1080/10635150600852011

Radies, D., Preusser, F., Matter, A., & Mange, M. (2004). Eustatic and climatic controls on the development of the Wahiba Sand Sea, Sultanate of Oman. Sedimentology, 51(6), 1359–1385. doi: 10.1111/j.1365-3091.2004.00678.x

Rambaut, A., & Drummond, A. J. (2013). Tracer v1. 5 Available from http://beast. Bio. Ed. Ac. Uk/Tracer. Accessed.

Rannala, B., & Yang, Z. (2003). Bayes Estimation of Species Divergence Times and Ancestral Population Sizes Using DNA Sequences From Multiple Loci. Genetics, 164(4), 1645–1656. doi: 10.1093/GENETICS/164.4.1645

Rannala, B., & Yang, Z. (2013). Improved Reversible Jump Algorithms for Bayesian Species Delimitation. Genetics, 194(1), 245–253. doi: 10.1534/genetics.112.149039

Riaño, G., Fontsere, C., Manuel, M. de, Talavera, A., Burriel-Carranza, B., Tejero-Cicuéndez, H., … Carranza, S. (2022). Genomics reveals introgression and purging of deleterious mutations in the Arabian leopard (*Panthera pardus nimr*). p. 2022.11.08.515636. bioRxiv. doi: 10.1101/2022.11.08.515636

Shults, P., Hopken, M., Eyer, P. A., Blumenfeld, A., Mateos, M., Cohnstaedt, L. W., & Vargo, E. L. (2022). Species delimitation and mitonuclear discordance within a species complex of biting midges. Scientific Reports 2022 12:1, 12(1), 1–13. doi: 10.1038/s41598-022-05856-x

Simó-Riudalbas, M., Pous, P. de, Els, J., Jayasinghe, S., Péntek-Zakar, E., Wilms, T., … Carranza, S. (2017). Cryptic diversity in *Ptyodactylus* (reptilia: Gekkonidae) from the northern hajar mountains of Oman and the United Arab Emirates uncovered by an integrative taxonomic approach. PLoS ONE, 12(8), 1–25. doi: 10.1371/journal.pone.0180397

Simó-Riudalbas, M., Tarroso, P., Papenfuss, T., Al-Sariri, T., & Carranza, S. (2018). Systematics, biogeography and evolution of *Asaccus gallagheri* (Squamata, Phyllodactylidae) with the description of a new endemic species from Oman. Systematics and Biodiversity, 16(4), 323–339. doi: 10.1080/14772000.2017.1403496

Šmíd, J., Sindaco, R., Shobrak, M., Busais, S., Tamar, K., Aghová, T., … Carranza, S. (2021). Diversity patterns and evolutionary history of Arabian squamates. Journal of Biogeography, 48(5), 1183–1199. doi: 10.1111/jbi.14070

Stange, M., Sánchez-Villagra, M. R., Salzburger, W., & Matschiner, M. (2018). Bayesian Divergence-Time Estimation with Genome-Wide Single-Nucleotide Polymorphism Data of Sea Catfishes (Ariidae) Supports Miocene Closure of the Panamanian Isthmus. Systematic Biology, 67(4), 681–699. doi: 10.1093/SYSBIO/SYY006

Sukumaran, J., & Knowles, L. L. (2017). Multispecies coalescent delimits structure, not species. Proceedings of the National Academy of Sciences, 114(7), 1607–1612. doi: 10.1073/PNAS.1607921114

Tamar, K., Chirio, L., Shobrak, M., Busais, S., & Carranza, S. (2019). Using multilocus approach to uncover cryptic diversity within *Pseudotrapelus* lizards from Saudi Arabia. Saudi Journal of Biological Sciences, 26(7), 1442–1449. doi: 10.1016/j.sjbs.2019.05.006

Tamar, K., Mitsi, P., & Carranza, S. (2019). Cryptic diversity revealed in the leaf-toed gecko *Asaccus montanus* (Squamata, Phyllodactylidae) from the Hajar Mountains of Arabia. Journal of Zoological Systematics and Evolutionary Research, 57(2), 369–382. doi: 10.1111/jzs.12258

Tejero-Cicuéndez, H., Patton, A. H., Caetano, D. S., Šmíd, J., Harmon, L. J., & Carranza, S. (2022). Reconstructing Squamate Biogeography in Afro-Arabia Reveals the Influence of a Complex and Dynamic Geologic Past. Systematic Biology, 71(2), 261–272. doi: 10.1093/SYSBIO/SYAB025

Thanou, E., Kornilios, P., Lymberakis, P., & Leaché, A. D. (2020). Genomic and mitochondrial evidence of ancient isolations and extreme introgression in the four-lined snake. Current Zoology, 66(1), 99–111. doi: 10.1093/cz/zoz018

Tonzo, V., Papadopoulou, A., & Ortego, J. (2019). Genomic data reveal deep genetic structure but no support for current taxonomic designation in a grasshopper species complex. Molecular Ecology, 28(17), 3869–3886. doi: 10.1111/mec.15189

Vences, M., Multzsch, M., Gippner, S., Miralles, A., Crottini, A., Gehring, P.-S., … Scherz, M. D. (2022). Integrative revision of the *Lygodactylus madagascariensis* group reveals an unexpected diversity of little brown geckos in Madagascar’s rainforest. Zootaxa, 5179(1), 1–61. doi: 10.11646/ZOOTAXA.5179.1.1

Vilaça, S. T., Piccinno, R., Rota-Stabelli, O., Gabrielli, M., Benazzo, A., Matschiner, M., … Bertorelle, G. (2021). Divergence and hybridization in sea turtles: Inferences from genome data show evidence of ancient gene flow between species. Molecular Ecology, 30(23), 6178–6192. doi: 10.1111/MEC.16113

Walters, A. D., Cannizzaro, A. G., Trujillo, D. A., & Berg, D. J. (2021). Addressing the Linnean shortfall in a cryptic species complex. Zoological Journal of the Linnean Society, 192(2), 277–305. doi: 10.1093/ZOOLINNEAN/ZLAA099

Wong, T. K., Cherryh, C., Rodrigo, A. G., Hahn, M. W., Minh, B. Q., & Lanfear, R. (2022, October 8). MAST: Phylogenetic Inference with Mixtures Across Sites and Trees (p. 2022.10.06.511210). p. 2022.10.06.511210. bioRxiv. doi: 10.1101/2022.10.06.511210

Yang, Z., & Rannala, B. (2010). Bayesian species delimitation using multilocus sequence data. Proceedings of the National Academy of Sciences, 107(20), 9264–9269. doi: 10.1073/pnas.0913022107

Zhang, C., Sayyari, E., & Mirarab, S. (2017). ASTRAL-III: Increased scalability and impacts of contracting low support branches. Lecture Notes in Computer Science (Including Subseries Lecture Notes in Artificial Intelligence and Lecture Notes in Bioinformatics), 10562 LNBI, 53–75. doi: 10.1007/978-3-319-67979-2_4/TABLES/3

